# High-throughput imaging of GABA fluorescence as a functional assay for variants in the neurodevelopmental gene, *SLC6A1*

**DOI:** 10.1101/2025.11.06.687003

**Authors:** Christopher Michael McGraw, Guoqi Zhang, Gillian Prinzing, Kimberly Wiltrout, Annapurna Poduri

## Abstract

Pathogenic variants in *SLC6A1*, which encodes GABA Transporter 1 (GAT-1), are associated with developmental delay, autism, epilepsy (e.g., epilepsy with myoclonic astatic seizures [EMAS]), and possibly schizophrenia. Functional assays to establish pathogenicity of human variants is a key limiting factor in the clinical interpretation of genetic findings. Methods based on radioactive [3H]-GABA uptake come with significant regulatory concerns, cost, and workflow complexity, which could be resolved by an alternate assay. To address this issue, we developed a high-content fluorescence imaging assay of GAT-1-mediated GABA uptake using a genetically encoded GABA sensor, iGABA-Snfr. We demonstrated that pathogenic variants strongly reduced uptake (mean, –89.4% [95% CI, –71.5% to –107.3%]). Some variants of uncertain significance (VUS) were associated with reduced GABA uptake (G111R, S459R, V511M; mean, –101.2% [95% CI –81.1% to –121.3%]), whereas others showed only mild reduction (R211C, R566H, F242V, R419C; mean, –33.6% [95% CI –17.2% to –50.1%]), supporting variant reclassification. Variant-specific effects on iGABA were highly correlated with the results of the radioactive [3H]-GABA assay (R^2^=0.8095, p<0.0001). The molecular chaperone 4-phenylbutyric acid (4PBA) was associated with ∼35% increase in iGABA. This non-radioactive assay is suitable for functional validation and high-throughput screening to identify positive modulators of GAT-1.

## 1. Introduction

The gamma-aminobutyric acid (GABA) transporter 1 (GAT-1), encoded by *SLC6A1*, plays a critical role in regulating extracellular GABA levels in the central nervous system (Richerson & Wu, 2004; Wu et al., 2007), thereby modulating inhibitory neurotransmission. Monoallelic variants in *SLC6A1* are associated with a spectrum of neurological disorders, including epilepsy, intellectual disability, autism spectrum disorder, and possibly schizophrenia (Cai et al., 2019; Carvill et al., 2015; Johannesen et al., 2018; Mattison et al., 2018; Silva et al., 2024). Most pathogenic variants have been shown to result in loss of function as evidenced by functional assays (Silva et al., 2024). The mechanism of GAT-1 haploinsufficiency involves reduced GABA transport due to variant-specific effects on GAT-1 activity (Mattison et al., 2018; Silva et al., 2024) as well as reduced GAT-1 surface expression (Cai et al., 2019; Nwosu et al., 2022; Silva et al., 2024) via protein destabilization and ER retention (Cai et al., 2019). Agonists of GAT-1 have not yet been identified. However, the molecular chaperone 4-phenylbutyric acid (4PBA) has shown therapeutic potential in S*lc6a1* mouse models (reduced spike-wave discharges) concomitant with increased GAT-1 surface expression and GABA uptake(Nwosu et al., 2022).

Functional assays are essential for evaluating the impact of variants on GAT-1 function and for reclassifying VUS to guide clinical management. Traditional methods for assessing GAT-1 function rely on radioactive [³H]-GABA uptake assays(Cai et al., 2019; Nwosu et al., 2022; Silva et al., 2024). While effective, radioactivity-based assays are burdened by regulatory restrictions and high waste disposal costs (Eglen & Reisine, 2009), making them less suitable for modern high-throughput applications such as high-content imaging.

To address these challenges, we developed a novel, non-radioactive, cell-based assay for GAT-1 function utilizing the genetically encoded GABA fluorescence sensor (iGABA-Snfr(Marvin et al., 2019) aka “iGABA”) coupled with high-throughput automated microscopy. The objectives of this study were two-fold: first, to validate the iGABA assay for assessing the functional impact of *SLC6A1* variants, including a novel missense variant M55T; and second, to demonstrate the assay’s suitability for high-throughput drug screening to identify positive modulators of GAT-1 activity. The GAT-1 iGABA assay provides a versatile platform for both variant classification and therapeutic development for SLC6A1-related disorders.

## 2. Materials and methods

### 2.1. Construction of hGAT1 and iGABA-Snfr expression vectors

To construct the hGAT1-P2A-tdTomato vector, the following sequences were amplified from the indicated sources by PCR: 1) full-length hGAT1 cDNA sequence without the final stop codon (1835 nts) from human clone (R&D Systems, hGAT-1/SLC6A1 VersaClone cDNA, RDC1033; Accession: NP_003033); 2) a 79-nt length P2A sequence from SID4x-dCas9-KRAB (AddGene 106399), and 3) tdTomato (1468 nts) from ptwB (AddGene 48735). Fragments were combined with the pCDNA3.1 vector (NeoR/KanR, AmpR) targeting the MCS by InFusion cloning (Takara) to generate an 8.6kB vector (➜ “V006-2”).

To construct an iGABA-Snfr-P2A-mBFP vector suitable for a cell-based assay, the following sequences were amplified from the indicated sources by PCR: 1) the CDS of iGABA-Snfr-F102Y-Y137L without the final stop codon (1976 nts) was amplified from pAAV-GFAP-iGABASnFR_F102Y_Y137L (Addgene #112174); 2) a 796 nt fragment containing P2A-mTagBFP2 was amplified from lentiGuide-Hygro-mTagBFP2 (AddGene# 99374). Fragments were combined with the pCDNA3.1 vector targeting the MCS by InFusion cloning to generate a 7.9kB vector (➜ “V007-2”). Subsequently, to abrogate surface expression, the 5’ IgG K-leader site was removed (InFusion) while retaining the start codon, and the 3’ terminal PDGFR site was removed, to yield a 7.8kB vector pCDNA3.1_iGABA-Snfr-F102Y-Y137L del IgGK del PDGFR T2A mBFP (➜ “V007-2.2”). Final vectors were Sanger sequenced through the coding sequences.

### 2.2. Generation of hGAT1 deletion constructs and human variant hGAT1 constructs

For deletion constructs, three sets of primers were designed to remove the extracellular domain (ECD; amino acids 145-211), cytoplasmic domain (CPD; amino acids 1-52), and transmembrane domains 6-7 (TM6-7, amino acids 292-342) based on Uniprot P30531. Representative human variants affecting the GAT-1 coding sequence were identified from ClinVar, gnomad, selected cohort studies(Carvill et al., 2015; Johannesen et al., 2018; Mattison et al., 2018), and the SLC6A1-Portal (https://slc6a1-portal.broadinstitute.org<u>/</u>, downloaded September 2022). Variants were selected based on recurrence across individuals and distribution across predicted pathogenicity, including variants classified as pathogenic/likely pathogenic (N= 7), benign/likely benign (N=2), and uncertain significance (N= 7). Clinical details were downloaded on September 8, 2022.

All variants are presented in **Table 1**. All primer sequences used for mutagenesis are provided in **Supplementary Table 1**. Deletions and variants were introduced into GAT-1 vector V006-2 by site-directed mutagenesis (InFusion Snap Assembly, Takara) and KOD polymerase (Sigma, 71085) using the associated primers (IDT). Constructs were verified by Sanger sequencing.

**Table 1.** Clinical and molecular characterization of GAT-1 variants investigated in this study.

| Human variant |  | Variant name (dbSNP) | Consequence specific allele | Protein allele | Variant start (aa) | PolyPhen2 prediction | PolyPhen2 | SIFT prediction | SIFT score | Variant supporting evidence | ClinVar or ACMG classification | Unique cases | % variant a/w epilepsy | iGABA fluorescence (%WT) |
| --- | --- | --- | --- | --- | --- | --- | --- | --- | --- | --- | --- | --- | --- | --- |
| p.M55T | Met55Thr | rs2124905468 | T/C | M/T | 55 | benign | 0.22 | deleterious | 0 | Phenotype_or_Disease, gnomAD | Uncertain significance | 2 | 50% (1/2) | +10% |
| p.G75R | Gly75Arg | rs1064795852 | G/A | G/R | 75 | possibly damaging | 0.931 | tolerated | 0.05 | Cited, Phenotype_or_Disease | Likely pathogenic / pathogenic | 3 | 100% (3/3) | -103.9% |
| p.G111R | Gly111Arg | rs1559622516 | G/A | G/R | 111 | probably damaging | 1 | deleterious | 0 | Phenotype_or_Disease | Conflicting interpretations (uncertain significance, likely pathogenic) | 2 | 50% (1/2) | -92.5% |
| p.R195H | Arg195His | rs773445048 | G/A | R/H | 195 | possibly damaging | 0.682 | tolerated | 0.07 | ExAC,Frequency, gnomAD, Phenotype_or_Disease, TOPMed | Benign | 1 | 100% (1/1) | +1.3% |
| p.R211C | Arg211Cys | rs756927822 | C/T | R/C | 211 | probably damaging | 1 | deleterious | 0 | ExAC,Frequency, gnomAD | Uncertain significance | 1 | 0% (0/1) | -36.8% |
| p.G232V | Gly232Val | rs1697444014 | G/T | G/V | 232 | probably damaging | 1 | deleterious | 0 | Phenotype_or_Disease | Likely pathogenic | 4 | 75% (3/4) | -108% |
| p.F242V | Phe242Val | rs1399297934 | T/G | F/V | 242 | benign | 0.002 | tolerated | 0.1 | Frequency,gnomAD, TOPMed | Uncertain significance | 2 | 100% (2/2) | -30.6% |
| p.A288V | Ala288Val | rs794726860 | C/T | A/V | 288 | probably damaging | 1 | deleterious | 0 | Cited, Phenotype_or_Disease | Likely pathogenic / pathogenic | 7 | 71.4% (5/7) | -93.6% |
| p.S295L | Ser295Leu | rs1574907241 | C/T | S/L | 295 | probably damaging | 1 | deleterious | 0 | Phenotype_or_Disease | Likely pathogenic* | 1 | 0% (0/1) | -103.3% |
| p.I322M | Ile322Met | rs757627416 | C/G | I/M | 322 | probably damaging | 1 | tolerated | 0.33 | ExAC,Frequency, gnomAD, TOPMed, Phenotype_or_Disease | Likely benign**, uncertain significance | 1 | 100% (1/1) | -31.6% |
| p.V342M | Val342M | rs760836450 | G/A | V/M | 342 | probably damaging | 0.999 | deleterious | 0 | Cited,ExAC,Frequency, gnomAD, Phenotype_or_Disease, TOPMed | Pathogenic | 11 | 90.9% (10/11) | -61.3% |
| p.A357V | Ala357Val | rs1553689859 | C/T | A/V | 357 | probably damaging | 0.997 | deleterious | 0.02 | Cited, Phenotype_or_Disease | Pathogenic | 4 | 100% (4/4) | -89.8% |
| p.G362R | Gly362Arg | rs1131691302 | G/A | G/R | 362 | probably damaging | 0.987 | deleterious | 0 | Cited, Phenotype_or_Disease | Likely pathogenic / pathogenic | 3 | 100% (3/3) | -79.7% |
| p.R419C | Arg419Cys | rs910130675 | C/T | R/C | 419 | possibly damaging | 0.682 | deleterious | 0.01 | Frequency,gnomAD, Phenotype_or_Disease, TOPMed | Uncertain significance | 1 | 100% (1/1) | -21.3% |
| p.S459R | Ser459Arg | rs1064795099 | C/A | S/R | 459 | probably damaging | 0.994 | deleterious | 0.02 | Cited, Phenotype_or_Disease | Conflicting interpretations (likely pathogenic, uncertain significance) | 2 | 100% (2/2) | -102.6% |
| p.V511M | Val511Met | rs1064794981 | G/A | V/M | 511 | probably damaging | 0.965 | tolerated | 0.06 | Cited, Phenotype_or_Disease | Conflicting interpretations (uncertain significance, likely pathogenic, pathogenic) | 3 | 100% (3/3) | -108.5 |
| p.R566H | Arg566His | rs767142926 | G/A | R/H | 566 | probably damaging | 0.998 | deleterious | 0 | ExAC,Frequency, gnomAD, Phenotype_or_Disease | Uncertain significance | 1 | 100% (1/1) | -45.9% |

Data related to number of unique cases and prevalence of epilepsy were retrieved from SLC6A1 Portal (Sept 8, 2022). iGABA fluorescence values are the average of two independent experiments.
\*, reported as "Uncertain Significance" in ClinVar, but "Likely pathogenic" supported by ACMG criteria due to PM1 (missense variant in mutational hot spot or well-defined domain), PP2 (missense variant in gene with low rate of benign missense variants), PM2 (extremely low frequency in gnomAD).
\*\*, the variant is not present in ClinVar, but predicted "Likely benign" in Ensembl Variant Effect Predictor. By ACMG criteria, it is consistent with VUS due to PM2 (low frequency in gnomAD), PP2 (missense variant in gene with low rate of benign missense variants)

### 2.3. Transient transfection assays of hGAT1 deletion and human variant constructs

HeLa cells (ATCC, CCL-2) were maintained in T75 flasks with EMEM (Cat#112-212-101, VWR Company) supplemented with 10% fetal bovine serum (FBS, Cat#16-000-04, Gibco) and 1X penicillin-streptomycin (PS) and 2% L-Glutamine (Cat# 16777-162, VWR) and incubated at 37°C with 5% CO2. For transient transfection, 2 million HeLa cells were seeded into each 60 mm petri dish on day 1. On day 2, for each dish, 2.5ug V007-2.2 and 2.5ug hGAT vector (WT or variant) (5ug total) were combined in 0.5mL OptiMEM and mixed gently. Lipofectamine 2000 (Cat#52758, Invitrogen) 10uL was diluted in 0.5mL OptiMEM. Diluted DNA and Lipofectamine were combined with gentle mixing and incubated 20 minutes at room temperature. Finally, the 1mL was added to each dish containing cells and 4mL of media, mixed gently by rocking back and forth, then incubated for 4-6 hours before replacing media with EMEM complete media. On day 3, transfected cells were dislodged by trypsin digest (Cat# 25300-054 from Gibco) and seeded at 500k cells per well into 96 well plate (Greiner, 655090). After 6 hrs of incubation, cells were washed twice with PBS and treated with 260 uM tiagabine (TGB; Cat # T3165, Lot PQ2JD-NP from TCI) diluted in EMEM + 10%FBS + 1X PS overnight (∼16 hrs). On day 4, media was removed and cells washed twice with PBS, then treated with gamma-aminobutyric acid (GABA) (Cat# A2129-100G, Sigma) diluted in EMEM at various concentrations at various incubations (1-8hrs) at 37degC. Each plate was subsequently imaged using the ImageXpress Micro (Molecular Devices).

### 2.4. Generation of hGAT1; iGABA-Snfr stable cell line

A stable HeLa cell line (ATCC, CCL-2) harboring both constructs (V006 and V007) was generated serially by transfection of linearized constructs (PvuI digest) followed by extended antibiotic selection, FACS to select single fluorescent cells for monoclonal culture, followed by FACS reassessment. A stable HeLa cell line was first established with V006-2 with G418 selection. This cell line was then used to generate a double stable cell line. The vector V007-2.2 was converted to exchange PuroR resistance cassette with the existing NeoR/Kan cassette by InFusion (➜ V007-2.3), but we failed to recover puromycin-resistant clones after several attempts. The vector V007-2.2 was then converted to have blasticidin resistance (➜ V007-2.4), from which 4 double-positive clones were successfully isolated by FACS, two of which grew well and retained double-fluorescence by FACS after extended incubation with double selection using blastocidin (1ug/mL) and G418 (200ug/mL). Clone #2 was used for all subsequent studies.

### 2.5. 4PBA assay

For testing 4-phenylbutyric acid (4PBA; Sigma P21005), the compound was dissolved to 1205.8mM in DMSO (198mg/mL) to generate a parent stock. GABA was prepared to 500mM (515.6mg in 2mL PBS) and further diluted to generate 50X GABA stocks for each GABA condition. 4PBA-GABA stocks for each condition were generated by diluting the parent 4PBA stock in EMEM (16.5uL into 6600uL EMEM) then combined with 50X GABA (161.7uL 4PBA + 3.3uL 50X GABA). For 4PBA 3mM, the parent 4PBA stock (1205.8mM) was used, but for 4PBA 1mM, the parent stock was first diluted to 400mM (DDW), then diluted in EMEM and combined with 50X GABA, as before. Drug plates were assembled in 96-well format using Integra Viaflow automatic multi-channel pipettors. On the day of testing, medium from prepared cells was aspirated and 1X 4PBA-GABA stocks were transferred directly to cells and incubated at 37degC for 6 hours prior to imaging.

### 2.6. Assays with hGAT1; iGABA-Snfr stable cell line

For assays using the hGAT1; iGABA-Snfr double stable cell line, cells were seeded into 384-well plates (5X10^4^ cells per well) in EMEM supplemented with 10% FBS, 2% L-Glutamine, 1X PS, G418 (200ug/mL), and blastocidin (1ug/mL) and allowed to grow to 80% confluence. On day 1, cells were preincubated overnight with 260uM TGB in EMEM. On day 2, GABA stock was diluted to 3X desired concentrations (10nm-5uM) in EMEM with 3X 0.1% DMSO on a “drug plate”, while EMEM was dispensed to an “EMEM only” plate using the Multidrop Combi dispenser (ThermoScientific). For GABA application, under semi-sterile conditions, the Bravo liquid handler (Agilent) was used to wash cells 3X in 1X PBS and then remove remaining PBS and apply 20uL from “DMEM only” and 10uL from “drug plate”. Plates were incubated at 37degC for 1-6hrs, before imaging using the IXM.

### 2.7. High-content imaging and automated analysis

Widefield fluorescence imaging was performed using the ImageXpress Micro (IXM) with 20X objective with the following filters and exposure durations: DAPI (mBFP; 10msec), FITC (iGABA-Snfr; 20msec), and TexasRed (tdTomato; 5msec). Image analysis was performed using Molecular Devices software to select cells with mBFP intensity above local background 150, with approximate width 5-50um. Per-cell average fluorescence values for mBFP, iGABA, and tdTomato are subsequently reported for all sites and wells.

### 2.8. In vitro assay of iGABA-Snfr fluorescence

In vitro fluorescence measurements were performed on a Tecan Spark plate reading fluorimeter at ∼28°C with excitation at 485 nm (20 nm bandpass) and emission collected at 532 nm (20 nm bandpass). Concentrated purified and dialyzed iGABASnFR.F102Y.Y137L protein was diluted to 0.2 μM in PBS. GABA and decoy compounds were purchased from Sigma-Aldrich and prepared as 100 mM stocks in PBS. Titrations were performed by making serial dilutions (1:2) of the stock compound into PBS and adding 10 μl of that to 100 μl of 0.2 μM protein solution. Fluorescence was measured before addition of compound, and ΔF/F was calculated as (F(treatment) − F(initial))/F(initial). Controls were performed with circularly permuted super-folder GFP (cpSFGFP; PMID: 16369541).

### 2.9. Data analysis and statistics

Data from IXM was extracted either as per-cell average fluorescence values or per-well averages across variable numbers of sites (4-10 sites) and processed further in R, Excel, and/or GraphPad Prism.

For variant experiments (with *k* variants), fractional change in iGABA fluorescence, *M*, due to GABA application is calculated in relation to each variant’s vehicle-treated control, as: *M_i_* = (F_i, GABA_ – F_i, vehicle_) / F_i, vehicle_, where F is the group-wise average iGABA fluorescence (across wells), and *i* is an element of the 1,…, *k* variants. Standard deviations, *s_i_*, were adjusted by: *s_i_*__adj_ = *s_i_ /* F_i, vehicle_. Normalization to WT was performed as: *M_i_adj_* = *M_i_ / M*_WT_, where *M_WT_* is the fractional change in iGABA fluorescence for wild-type GAT-1 control. For %WT, *M_i_adj%_* = *M_i_adj_* X 100.

For stable cell line experiments, fractional change in iGABA fluorescence, M, due to GABA application is calculated (similar to above) in relation to vehicle-treated control, but with *i* as an element of 1, …, *p* levels of GABA concentration.

Statistical inference was performed in GraphPad Prism as described in the main text. For plate normalization using linear models, per-well average data was analyzed in R using a custom script. Linear models were fit using function *lm* (R package, *stats*) while linear mixed models were fit using function *lmer* (R package, *lme4*). Bootstrap simulation was performed in R using the mean and standard deviations for each control. The robust strictly standardized mean difference (RSSMD) was calculated by the uniformly minimum-variance unbiased estimator (UMVUE) method(Zhang, 2011).

## 3. Results

### 3.1. Development of a cell-based assay for hGAT1 function using iGABA-Snfr fluorescence

To detect changes in intracellular GABA mediated by hGAT1 after exogenous application of GABA (**Fig. 1B**), we generated two vectors, one expressing human GAT-1 (hGAT1) cDNA (Accession: NP_003033) stoichiometrically linked with the red fluorescent fluorophore tdTomato (**Fig. 1A**) and another expressing modified iGABA-Snfr stoichiometrically linked to the blue fluorescent fluorophore mTagBFP2 (abbreviated mBFP). We selected the HeLa cell line (ATCC, CC2) based on publicly available RNA-seq data (2021 CCLE release; DepMap, ACH-001086) showing undetectable expression of *SLC6A1* (GAT-1; 0 TPM) and low or undetectable expression of other GABA transporters including *SLC6A13* (GAT-2; 0 TPM), *SLC6A11* (GAT-3; 0 TPM); *SLC6A12* (BGT1; 0 TPM), and *SLC6A6* (TauT; 1-2 TPM). Using automated high content fluorescence microscopy (ImageXpress Micro) and automated image analysis (**Fig. 1C)**, we demonstrated the ability to assay levels of iGABA-Snfr fluorescence from thousands of cells per condition in 96-well format with a significant increase in average per-cell iGABA fluorescence in response to exogenous GABA application (**Figure 1D-E).**

**Figure 1.**
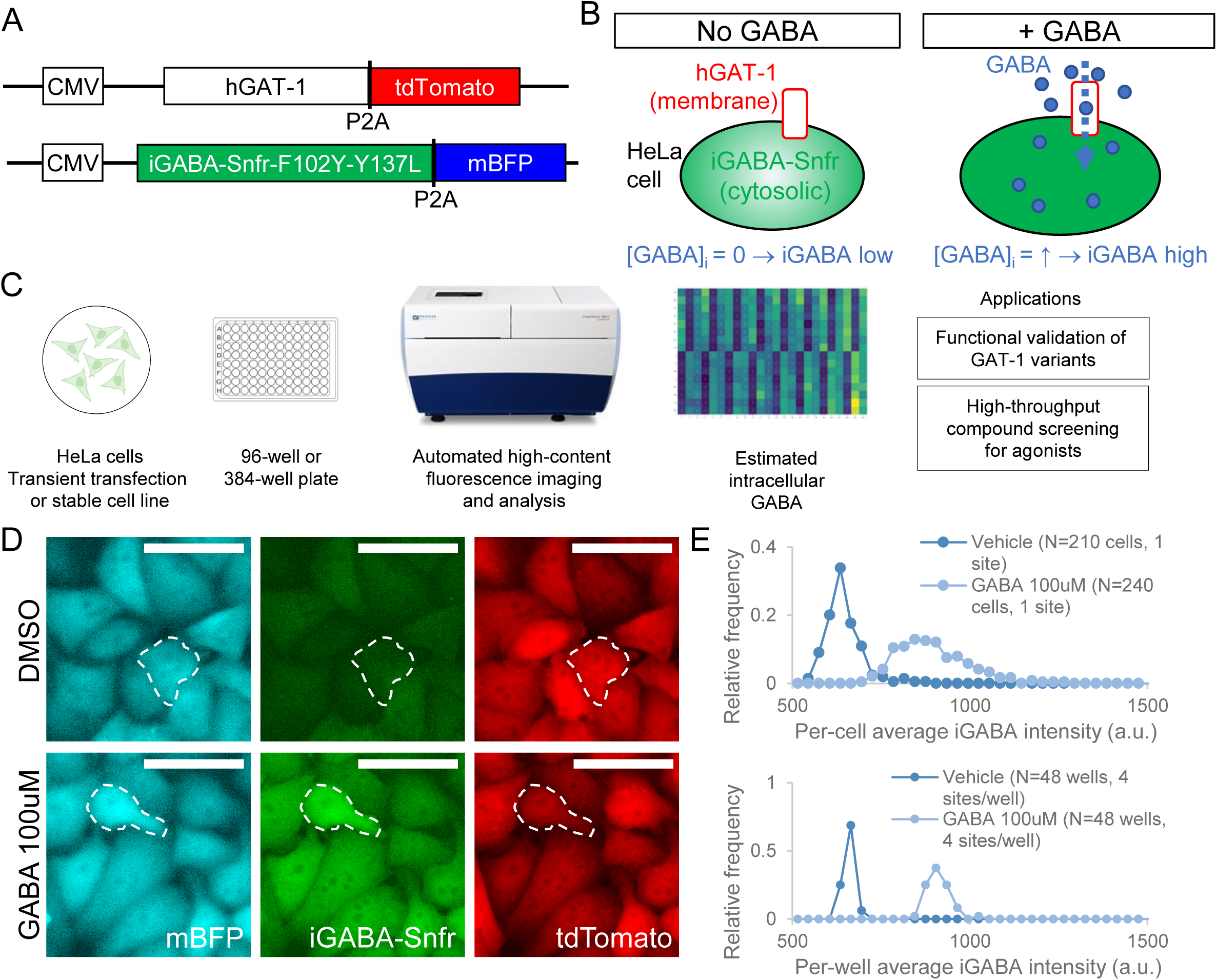
Development of a cell-based assay for hGAT-1 function using iGABA-Snfr fluorescence. (**A**) Schematic of constructs (pCDNA3.1) used to express human GAT-1 (*upper*) and a genetically encoded GABA sensor, iGABA-Snfr (*lower*) with stoichiometrically linked fluorophores. Not to scale. (**B**) Approach to quantifying GAT-1-mediated GABA uptake in HeLa cells. In the presence of exogenous GABA and GAT-1 protein, elevated intracellular concentration of GABA ([GABA]_i_) is reflected by increased fluorescence by iGABA-Snfr. (**C**) Workflow for assays to validate the function of GAT-1 variants and the detection of agonist compounds. (**D**) Representative images of iGABA assay and mock-up of automated analysis to measure per-cell average iGABA-Snfr fluorescence. Scale bar = 50 um. (**E**) Relative frequency histograms of iGABA-Snfr fluorescence after incubation with GABA (100uM) versus vehicle (DMSO), as measured by automated analysis at the per-cell level within a single site (*top*) and the per-well averages (*lower*) across a representative 96-well plate (N=48 wells per condition, 4 sites per well).

### 3.2. Validation of relative change in per-cell GABA fluorescence as molecular read-out of GABA transporter activity

To benchmark the effects of null variants, we utilized deletion constructs targeting domains previously demonstrated to be critical for GAT1-mediated GABA uptake (CPD, ECD, TM6-7) (**Fig 2A**). We confirmed that deletion variants were associated with significantly reduced iGABA fluorescence relative to wild-type across a 5-log range (1µM to 10mM; **Fig 2B**). Expressed as normalized fractional change in iGABA (relative to vehicle controls; **Fig 2C**) or normalized relative to wild-type (**Fig 2D**), the reduced uptake in GABA appeared most specific at lower concentrations of extracellular GABA (1-10uM, similar to GAT-1 Km ∼3uM(Wu et al., 2007)). At higher concentrations (100 µM GABA), we observed ∼30-40% of WT GABA uptake, which we hypothesized may be due either to residual GAT-1 protein function or nonspecific lower affinity mechanisms of uptake. To assess this question, we examined iGABA fluorescence in cells without GAT-1 transfection (“no GAT1” controls). Surprisingly, we observed significantly greater nonspecific GABA uptake in the absence of GAT-1 at both low (1uM) and higher (100uM) GABA concentrations compared to deletion variants (**Supplementary Fig 1**), suggesting active interference between GAT-1 deletion constructs and unknown mechanisms mediating low-affinity GABA uptake in HeLa cells (perhaps TauT, a low-affinity GABA transporter), despite undetectable GABA uptake by these variants. To further optimize the assay, we tested the effect of overnight pre-treatment with the GAT-1 selective inhibitor, tiagabine (TGB) followed by wash-out, and showed increased fractional change in iGABA across all GABA concentrations (**Supplementary Fig 2A**). We also tested a range of incubation durations and demonstrated increasing uptake up to 6 hours post-treatment (**Supplementary Fig 2B**). Based on these observations, we determined that TGB pretreatment followed by low GABA concentration (1 µM) incubation x 6 hrs would be most suitable for further experiments. We demonstrated that increased iGABA fluorescence under these conditions could be detected robustly across replicates (robust SSMD >2, robust Z prime >0.1; **Fig 2E**).

**Figure 2.**
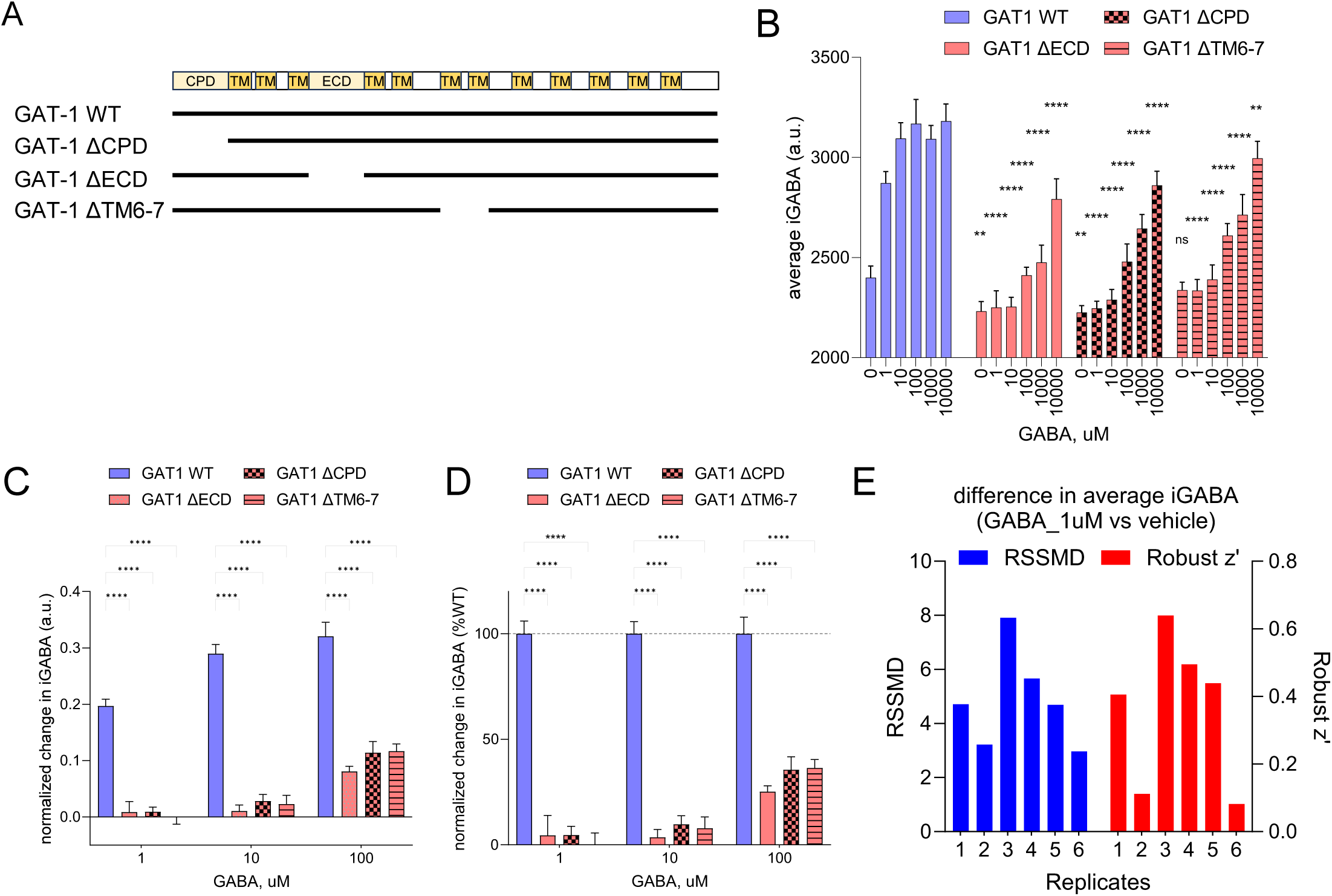
Validation of iGABA-Snfr assay for assessing GAT-1 loss-of-function variants. (**A**) Schematic of full-length human GAT-1 protein (Uniprot P30531) and the location/range of deletions implicated in ΔCPD (cytoplasmic domain), ΔECD (extracellular domain), and ΔTM6-7 (transmembrane domains 6 and 7). (**B**) GABA concentration series (5-log range) demonstrates enhanced intracellular GABA accumulation in wildtype GAT-1 compared to GAT-1 constructs harboring deletions of critical domains (ECD, CPD, and TM6-7). (Two-way ANOVA, with Dunnett’s multiple comparisons test; 6 families, 3 comparisons per family, relative to WT). (**C-D**) Normalized change in iGABA fluorescence (C) and as %WT (D) is significantly greater in wildtype GAT-1 versus GAT-1 deletion constructs at GABA concentrations 1uM-100uM. For each construct, data are the difference in iGABA fluorescence between GABA conditions and vehicle control, divided by vehicle control. (Two-way ANOVA, with Dunnett’s multiple comparisons test; 3 families, 3 comparisons per family, relative to WT). (**E**) The robust strictly standardized mean difference (RSSMD) and robust z-prime are calculated between positive (GABA 1uM) and negative (vehicle) controls (*right*) across replicate plates. For B-D, assay format is 96-well plate, transient transfection, incubation time 6-7 hrs. Data are the mean +/-SEM of independent per-well averages of iGABA (4 wells per condition (construct x GABA concentration), 4 sites per well)

### 3.3. GABA fluorescence provides a functional assessment of missense variants in GABA transporters

We next asked whether iGABA fluorescence could detect changes in GABA uptake related to human missense variants in *SLC6A1*. For this purpose, we selected *SLC6A1* variants associated with neurodevelopmental disorder and/or epilepsy (**Fig 3A**), including 7 pathogenic or likely pathogenic variants (G75R, G232V, A288V, S295L, V342M, A357V, G362R), two benign or likely benign variants (R195H, I322M), and 7 variants of uncertain significance (VUS;, G111R, R211C, F242V, R419C, S459R, V511M, R566H, S574R). One of these variants (R211C) has also been associated with schizophrenia(Rees et al., 2020). Together, these variants comprise 38.7% (n=12 of 31) of all recurrent missense variants to date (Silva et al., 2024).

**Figure 3.**
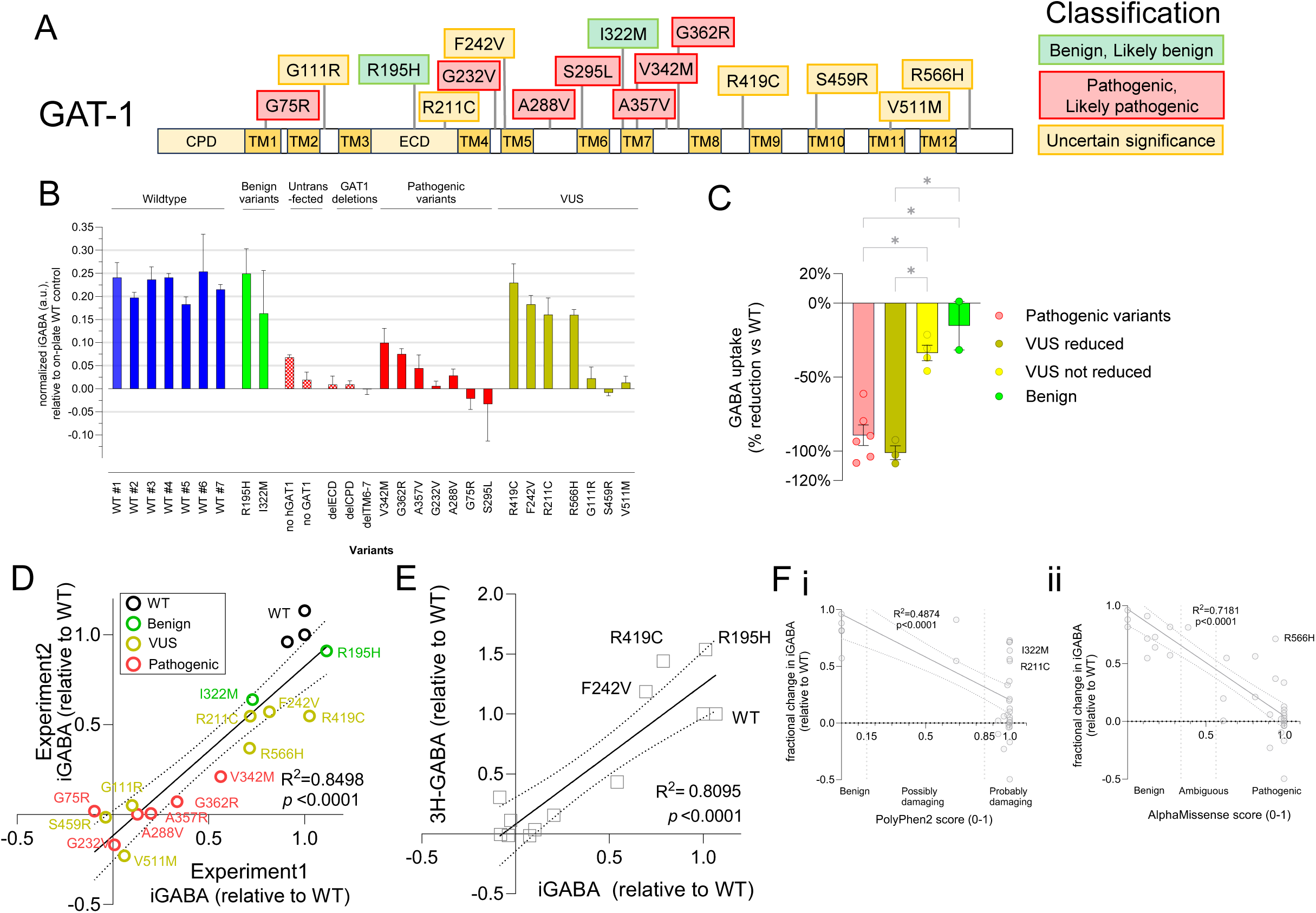
Validation of iGABA-Snfr assay for assessing human GAT-1 variants. (**A)** Schematic of human GAT-1 protein (Uniprot P30531) and human variants tested in this study. Boxes indicate the amino acid substitution associated with each missense variant. Color coding indicates clinical classification based on ClinVar. (**B)** Representative results of iGABA assay as a function of variant. Data are fractional changes in GABA fluorescence expressed as mean +/-sem in the presence of GABA 1uM relative to untreated control for each construct (4 wells per condition). (**C)** Significant impairments in fractional iGABA change is noted in all pathogenic variants tested and a subset of VUS. Data are mean +/-sem for grouped variants shown in B. (n=4 wells per condition). Kruskal-Wallis test with Dunnett’s test; *\*, p*<0.05. (**D)** iGABA test-retest correlation for GAT-1 variants is high (R^2^=0.8498). Data are fractional GABA uptake relative to WT control from two independent experiments (n=4 wells per condition). (**E**) Variant-specific GABA fluorescence is closely correlated (R^2^=0.8095) with published 3H-GABA uptake assays. Data is adapted from the “UCSF dataset” published by Silva et al AJHG 2024 (https://doi.org/10.1016/j.ajhg.2024.04.02). (**F)** Significant correlations between fractional iGABA and PolyPhen2 (i) and AlphaMissense (ii) scores

We introduced missense variants into hGAT1 by site-directed mutagenesis and performed the iGABA assay following transient transfection. We observed consistent reductions in normalized GABA uptake among pathogenic variants (**Fig 3B-C**) with – 89.4% mean reduction in iGABA (SD 17.1, 95% CI –71.5% to –107.3%; N=6). Using the lower limit (95^th^ CI) of pathogenic GABA uptake as a threshold, some VUS were associated with reduced GABA uptake (G111R, S459R, V511M; mean reduction, – 101.2%, SD 8.10, 95% CI –81.1% to –121.3%, N=3), whereas others showed only mild reduction (R211C, R566H, F242V, R419C; mean reduction, –33.6%, SD 10.4, 95% CI – 17.2% to –50.1%, N=4) (**Fig 3C**).

Test-retest analysis demonstrated that variant-specific iGABA was highly correlated (R^2^=0.8498, *p*<0.0001) between independent replicates (**Fig 3D**). Variant-specific effects on iGABA are also highly correlated (R^2^=0.65-0.81, *p*<0.0001) with previously published measurements estimated by radioactive [^3^H]-GABA transport assay(Silva et al., 2024)(**Fig 3E**), suggesting that the iGABA method performs similarly to this gold-standard method. Last, variant-specific reductions in iGABA correlate with several pathogenicity prediction scores, including PolyPhen2 (R^2^=0.49, p<0.0001; **Fig 3F.i**), SIFT (R^2^=0.42, p<0.0001; not shown), and AlphaMissense (R^2^=0.7181, p<0.0001; **Fig 3F.ii**).

Radioactive assays estimate an instantaneous rate of GABA uptake (pmol/ug/min) whereas the iGABA method estimates intracellular GABA as it approaches equilibrium, reflecting the balance between rates of GABA uptake vs GABA removal due to unspecified processes (metabolism, sequestration, efflux). To understand these differences, we performed time-series analysis to assess whether a pathogenic variant possessing some residual transporter activity (V342M) alters the rate of GABA uptake, the steady state intracellular concentration of GABA, or both (**Fig 4A**). Based on a non-linear fit of pseudo-first order kinetics, the pathogenic V342M variant reduced steady-state GABA (Plateau: WT: 0.28, 95% CI 0.26-0.31, vs V342M: 0.11, 95% CI 0.087-0.204; **Fig 4B**) but had no effect on the rate at which iGABA approached its variant-specific steady-state (rate constant, K (min^-1^): WT: 0.0126, 95% CI 0.008-0.020; V342M: 0.009, 95% CI 0.002-0.018; **Fig 4C**). This may explain the high correlation that we observed between iGABA and the rates of GABA uptake estimated by radioactive methods.

**Figure 4.**
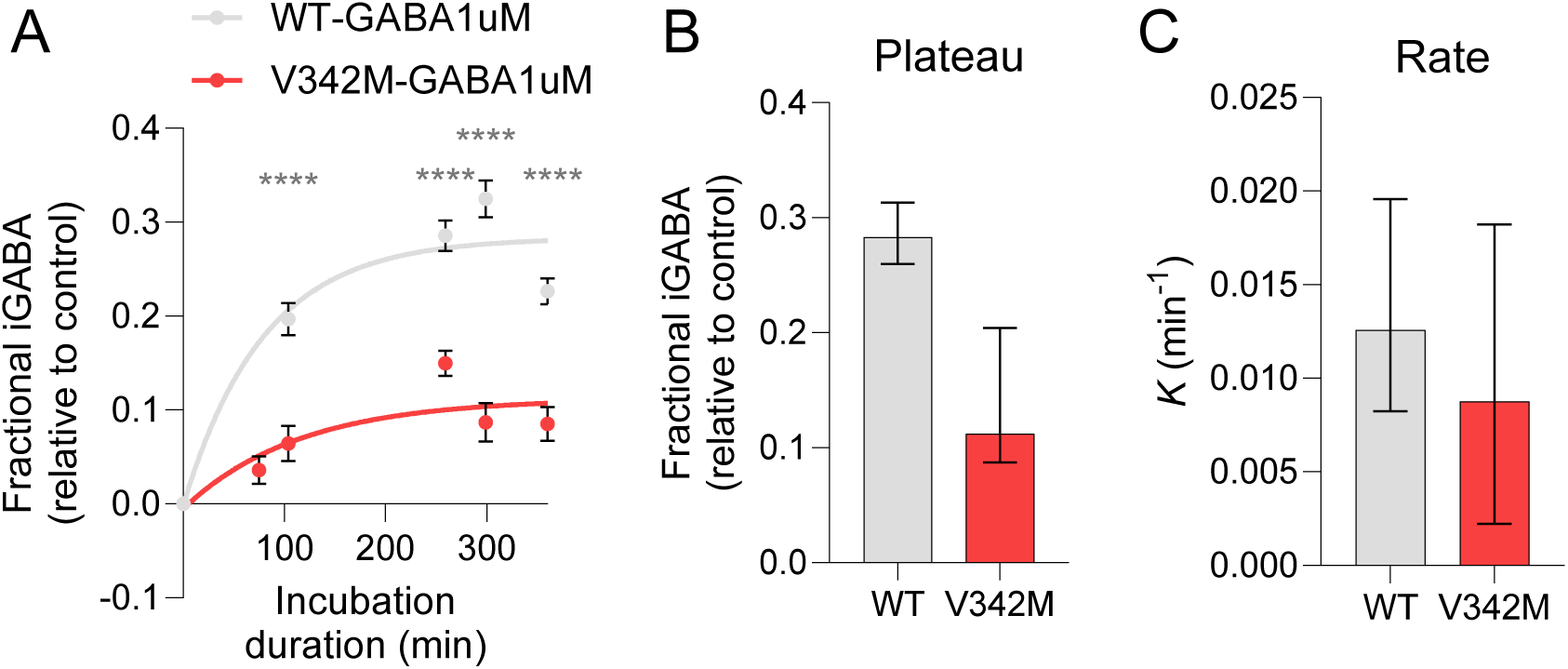
Pathogenic variants in GAT-1 are associated with reduced steady-state iGABA. (**A**) Time-course experiments following 1-6 hours incubation demonstrate marked reduction in plateau of [GABA]_in_ mediated by mutant GAT-1 V342M versus GAT-1 WT control. Data are average +/-s.e.m. (n=15 wells per timepoint x condition). Non-linear fits were computed based on pseudo-first order association kinetics, according to the equation `Y=Y0 + (Plateau-Y0)*(1-exp(–K*x))`. Statistics are unpaired t-test with Welch correction with FDR correction. ****, adjusted *p*<0.0001. (**B-C**) Parameters from non-linear fit of (A) shows significantly reduced steady-state levels (parameter Plateau; B) but no significant difference in the rate constant (parameter K; C). Data are best-fit parameter values +/-95% C.I.

### 3.4. GABA fluorescence may model haploinsufficiency of GAT-1 loss-of-function variants in the context of wild-type GAT-1 co-expression

As most variants reported in association with disease are monoallelic(Silva et al., 2024), we next sought to assess the effects of combining WT with mutant hGAT1 variants in the iGABA assay, mimicking the heterozygous state. We observed that the 1:1 combination of WT GAT-1 with pathogenic GAT-1 variants routinely increased measured GABA uptake to a level ∼50% 2xWT (**Fig 5A**). Meanwhile the benign I322M variant with partially reduced iGABA increased to ∼100% 2XWT (**Fig 5A**). Both findings are expected for a simple additive effect and consistent with the putative disease mechanism of haploinsufficiency. This also suggests that the transfection quantities used for the assay were not saturating and that the read-out is in a linear range.

**Figure 5.**
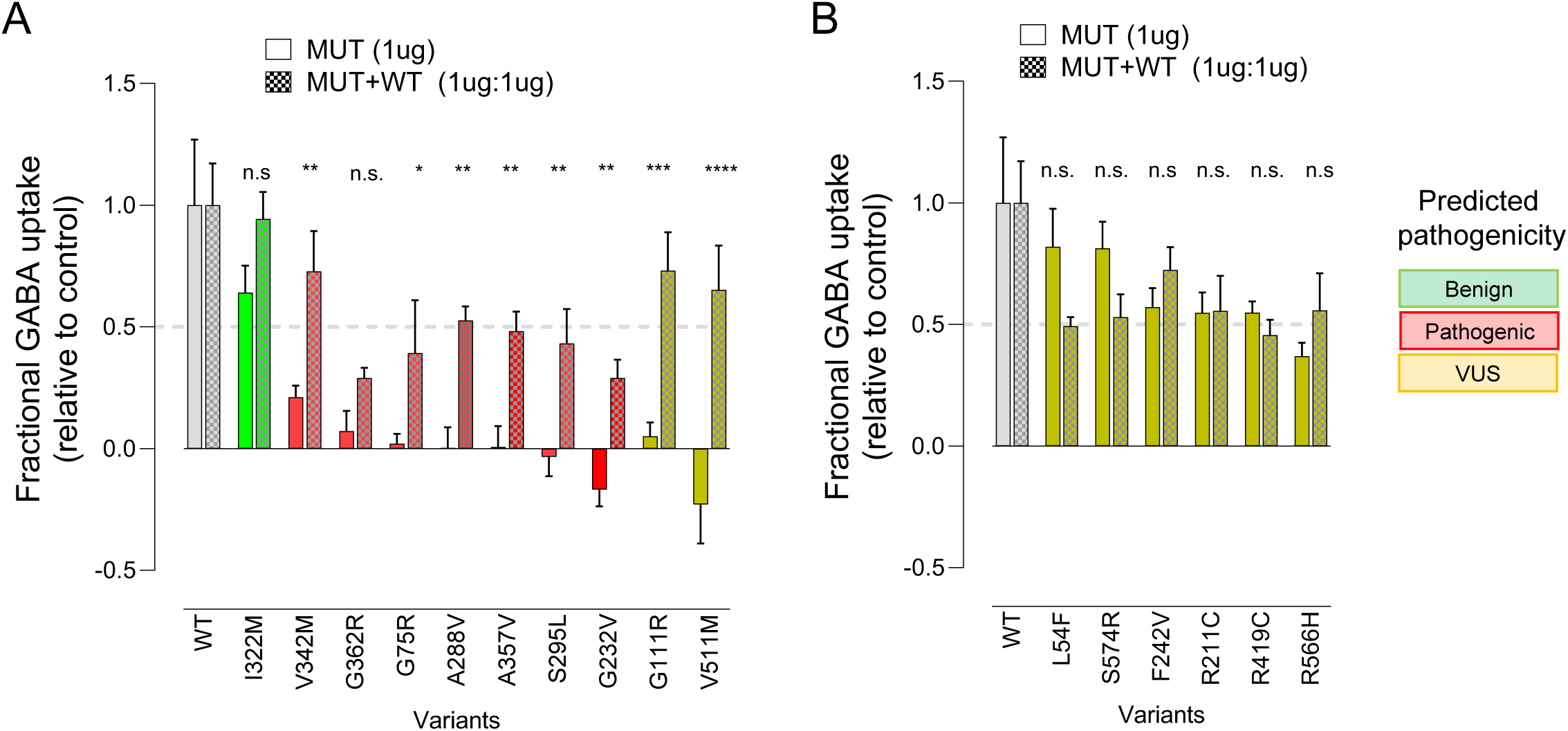
Complex variant-dependent interactions between WT and mutant GAT-1 affect GABA uptake. (**A-B**) Effects of combining GAT-1 variants with equimolar WT GAT-1 by transient transfection show additive (A) and non-additive (B) effects. Data are fractional change in iGABA relative to control (4 wells per condition), presented as mean +/-sem. The MUT series (*open boxes*) are normalized by WT, while MUT+WT data (*checkerboard*) are normalized by 2xWT. Statistics are Dunnett’s test for differences between MUT vs MUT+WT for each variant followed by false-discovery rate (FDR) correction (n.s., not significant. *\*, adj p*<0.05. **, *adj p*<0.01. ***, *adj p*<0.001).

**Figure 6.**
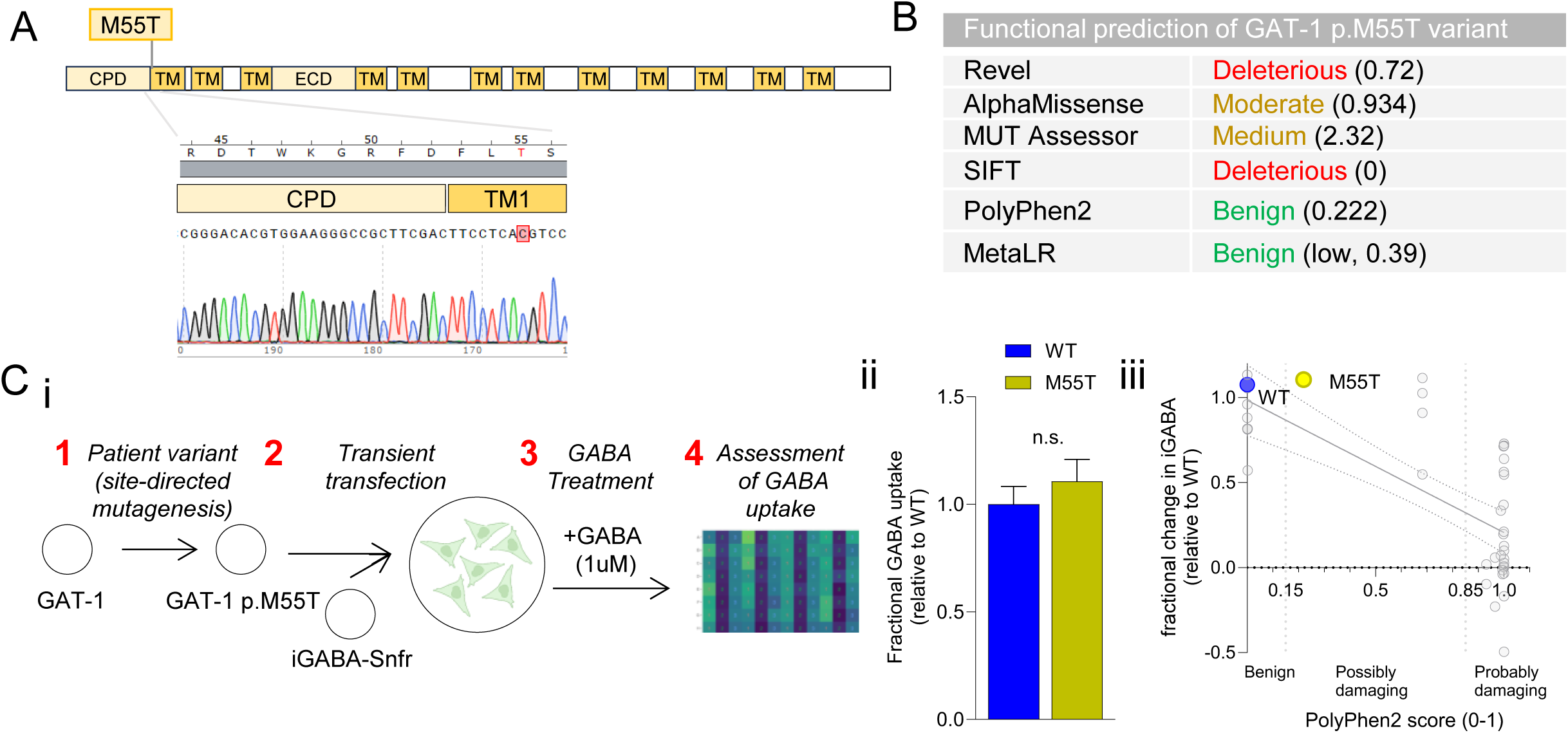
iGABA assay can evaluate functional effects of novel coding sequence variants on GABA transport. (**A)** The p.M55T variant is located within the 1^st^ transmembrane (TM) domain of GAT-1 (Uniprot P30531). Sanger sequencing of the GAT-1 p.M55T construct used in iGABA assay is shown. (**B)** Pathogenicity prediction scores for p.M55T are equivocal, with several suggesting deleterious effect but others not (PolyPhen2, MetaLR). **(C**) Approach to assay the effects of novel human GAT-1 variants on iGABA (i) and its negligible effects on GABA uptake (ii), in good correspondence with PolyPhen2 score (iii). Data are mean +/-sem, expressed relative to WT controls (n=15 wells per condition; 6 sites per well). Statistics are unpaired T-test with Welch’s correction. n.s., not significant.

For two VUS associated with reduced GABA uptake (G111R, V511M) by the iGABA method, we also observed an additive effect when combined with WT GAT-1 (**Fig 5A**). However, for several VUS associated with only mild reductions in iGABA (L54F, R211C, F242V, R419C, R566H, S574R), the addition of WT GAT-1 did not have the same additive effect (6 of 6 VUS with reduced iGABA). This unexpected finding may suggest an artifact of the heterologous system, but that is difficult to reconcile with the results observed for pathogenic variants, the majority of which yielded evidence of an additive effect (5 of 6 variants tested). Alternatively, these findings may suggest a disease mechanism acting against the compensatory effects of WT GAT-1.

### 3.5. GABA fluorescence provides insights into novel missense variant in GAT-1

We used the iGABA assay to assess the effect of a novel missense variant in *SLC6A1* in a patient with non-acquired focal epilepsy (NAFE).

Briefly, the proband was born at term with perinatal history complicated by intrauterine drug exposure and planned repeat Cesarean section under general anesthesia. At delivery, Apgar scores were 4 and 0 at one and five minutes, respectively, with intubation in the delivery room and extubation to room air on day of life 1. He walked independently by 15 months of age. However, language and social development were delayed. He was diagnosed with autism spectrum disorder at 3 years old. At age 3 years, he spoke in word approximations and echolalia. He was quick to anger and displayed head-banging and aggressive behavior. He made great progress in language and social development between the ages of 3.5-4 years in a preschool program for children with autism spectrum disorder. At 6 years old, he could participate in simple games, write and name some letters and numbers, cut with scissors, and use a brush to make paint strokes. He toilet-independently by 5 years old.

His seizure onset was at age 4 years, initially with semiology of staring with behavioral arrest, and he later developed seizures with left hemiclonic semiology. He has had 3 lifetime generalized tonic-clonic seizures. His seizure frequency increased over time and he experienced a developmental regression at 6 years old when seizures were poorly controlled. He lost the previously acquired language skills, interest in socializing, his behavior had worsened, and his gait became clumsy and somewhat crouched. At current age of 12 years, he has not regained the skills he lost despite improved seizure control. At last exam at 12 years old, he was able to speak in short sentences with quiet speech intermixed with jargoning and echolalia. His initial EEG showed right posterior temporal occipital epileptiform spikes. Subsequent EEGs at ages 6-9 years showed frequent sleep-potentiated multifocal spikes. At 11 years of age, an EEG showed runs of temporal-parietal focal spikes in sleep. He currently takes lamotrigine, which has been effective at seizure control. In the past, Prior regimens have included levetiracetam (stopped due to behavioral side effects), oxcarbazepine (ineffective), and valproic acid (the most effective but limited use due to side effect of thrombocytopenia). He also has a significant tremor in bilateral upper extremities that worsened with valproic acid but did not resolve after valproic acid was discontinued. He also continues to have tics, moderate intellectual disability, cortical visual impairment, behavioral problems attributed to ADHD and mood dysregulation, as well as severe constipation. For his behavioral problems, which include impulsive behavior and aggression toward self and others, he has trialed multiple formulations of stimulant medications and antipsychotic medications. Behaviors have improved the most with guanfacine and valproic acid.

Radiological testing including MRI brain was unremarkable including no evidence of hypomyelination. He had unremarkable chromosomal microarray (CMA) and Fragile X testing. Whole-exome sequencing, updated in 2024, identified a variant in *SLC6A1*, c.164 T>C (chr3-11059061; dbSNP, rs2124905468; GRCh38.p14 NC_000003.12, exon 3 of 16) resulting in p.M55T (Refseq, NM_003042.4). This variant is located in the 1^st^ transmembrane domain (**Fig 4A**), which is a recognized mutation hotspot(Silva et al., 2024). It is predicted to be pathogenic by several pathogenicity prediction scores (Revel, AlphaMissense, MUTAssessor, SIFT) though others classify it as benign (PolyPhen2; MetaLR) (**Fig. 4B**). The variant is not present in the general population (gnomAD) and is reported once in ClinVar as uncertain significance with limited clinical description. Parental DNA was not available for segregation analysis as the patient was adopted. Taken together, the patient’s clinical presentation with seizures, tremor, and intellectual disability was believed to be reasonably well-explained by the *SLC6A1* p.M55T variant but with an ACMG classification of uncertain significance (criteria, PM2, PP3).

The patient also harbored a heterozygous missense variant in GRIK2 c.1744G>A, p.A582T (chr6-102337734; dbSNP, rs2484646253; GRCh38.p14, NM_021956.5, exon 12 of 17), which was not reported in ClinVar or other literature and was interpreted as having uncertain significance. The VUS in GRIK2 was believed less likely to be a primary cause of patient’s epilepsy (criteria PM2, PP3) but cannot be fully excluded. GRIK2 has recently been associated with neurodevelopmental disorders with DD and prominent gait abnormalities(Stolz et al., 2021). The most severely affected patients (3 children with de novo variants at p.T660K and 2 children with p.T660R variants) presented with global developmental delay, non-ambulatory status, onset of epilepsy before age 3 years (4 out of 5 within 1^st^ 12 mos), and brain MRI abnormalities in the first years of life suggesting hypomelination, while the least severely affected patients (4 children and 1 adult with variants at p.A657T) presented with ID, DD, and incoordination, but did not develop epilepsy and had normal MRIs(Stolz et al., 2021).

The reported variants cluster in two critical functional domains, the pore-forming M3 transmembrane helix and the adjacent M3-S2-gating linker(Stolz et al., 2021). In contrast, our patient’s GRIK2 variant (p.A582T) is not located in the implicated domains, he developed seizures at age 4 yrs, ambulates, and has normal brain MRI, suggesting poor overlap with known cases of GRIK2-associated NDD. To understand the functional effects of the *SLC6A1* p.M55T variant, we assessed its effects on GAT1-mediated GABA uptake using the iGABA assay (**Fig 4C.i**).

Surprisingly, the M55T variant is not associated with detectable reduction in GABA uptake compared to WT controls (WT: 1 (mean) +/-0.083 (sem), M55T: 1.11 +/-0.102; *p*=0.428, Welch’s unpaired t-test; **Fig 4C.ii**), consistent with the PolyPhen2 prediction (**Fig 4C.iii**). These findings may suggest limitations of GABA uptake assays in heterologous cell lines, or they may suggest that the *SLC6A1* p.M55T variant does not account for this patient’s epilepsy.

### 3.6. Assessment of iGABA reporter for drug responses and high-throughput screening

We hypothesized that the iGABA-Snfr fluorescence assay of GAT-1 activity could be suitable for screening compounds to identify new modulators of GABA transporter activity.

To test this hypothesis, we developed a double stable cell line expressing both iGABA-Snfr and hGAT1 constructs (Methods), and used the iGABA assay to assess the response to the candidate compound, 4-phenylbutyric acid (4PBA) (**Fig 7A**). 4PBA was previously described to have positive effects on GAT-1 through increased surface expression of mutant and WT GAT-1(Nwosu et al., 2022). Given our previous findings showing increased steady state iGABA in response to elevated hGAT1 dosage, the correlation between rate of GABA uptake and steady state iGABA and the effect of pathogenic variants on these parameters, we predicted that if 4PBA is therapeutically effective, we should be able to detect its effect in our assay. Indeed, we observed a significant ∼35% increase in iGABA after treatment with 4PBA (1-3mM) in the setting of GABA 1uM (vehicle: 0.434 (mean) +/-0.006 (sem) fractional change in iGABA, vs 4PBA_1mM: 0.58 +/-0.015, *p*= 0.0057; vs 4PBA_3mM: 0.595 +/-0.020, *p*= 0.0023, Dunnett’s multiple comparison test) after 6 hour co-incubation (**Fig 7B**). These effects were not significant at lower GABA (100nm). At higher GABA (10uM), the effect of 4PBA was equivocal, with lower concentration (4PBA_1mM; *p*= 0.0112) and a DMSO control (*p*=0.0288) both significantly different from Vehicle1, but not higher concentration (4PBA_3mM; *p*=0.195). This effect is not likely to be mediated by direct interactions between 4PBA and the iGABA-Snfr sensor, as no increase in iGABA was detected by direct *in vitro* testing (**Supplementary Fig 3**). These findings suggest that the effect of 4PBA is detectable under specific assay conditions, despite differences with previously reported assays(Nwosu et al., 2022), further corroborating its utility in drug discovery.

**Figure 7.**
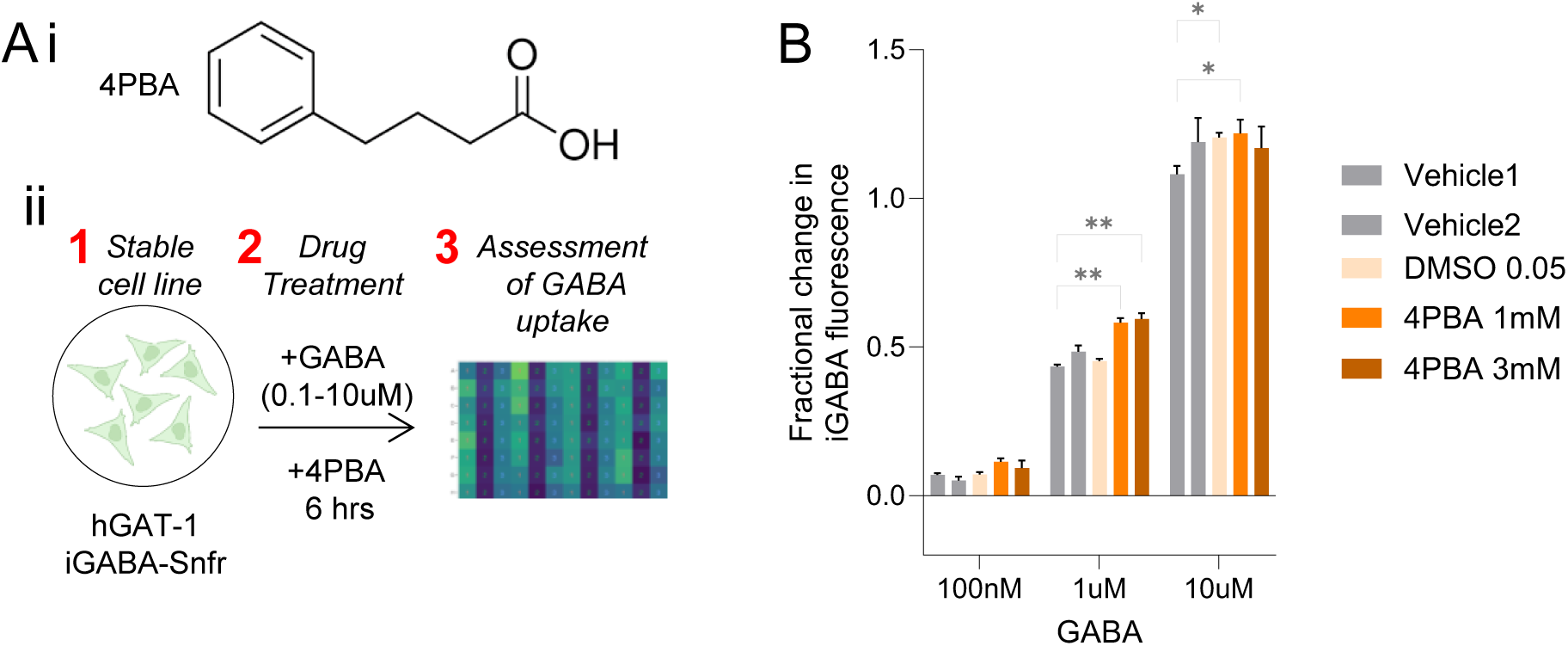
Mild effect of 4-phenylbutyric acid (4PBA) on GAT-1 activity in iGABA assay. (**A**) Structure of 4-phenylbutyric acid, 4PBA (i) and schematic for experiment (ii). (**B**) Mild enhancement of iGABA by 4PBA following application of GABA (0.1-10uM). Data are mean +/-sem. (n=4 wells per condition; 96-well format; 6 hour incubation). Controls include two duplicate vehicle controls (H2O; Vehicle1-2), and a DMSO (0.05%) control. Statistics are one-way ANOVA followed by Dunnett’s multiple comparison relative to control (“Vehicle 1”). n.s., not significant. *, *p*<0.05, **, *p*<0.01, ***, *p*<0.001.

We next evaluated the performance of the iGABA assay using the stable cell line in 384 well format using automated liquid handling devices and the high content fluorescence imager (**Fig 8A**). To understand the sources of assay variance, we performed preliminary testing with a 4-log range of GABA (10nM, 100nM, 1uM, 10uM) across 2×384 well plates, and fit covariates using a linear model (**Fig 8B.i**). We observed that ∼88.2% variance in iGABA was explained by this model (residual R^2^, 0.1178), with the GABA concentration accounting for the majority (71.4%, *p*=2.20E-16) followed by intermediate contributions from the Column position (13.3%, *p*=2.20E-16), and minimal contributions by Column*Order (1.3%, *p*=1.64E-15), Plate (1.1%, *p*=3.33E-13), Row (0.7%, *p*=6.13E-09) and Row*Column (0.4%, *p*=8.52E-06). These findings suggested that while GABA dose drives fluorescence largely independently of plate layout or minor differences in incubation duration introduced by serial imaging, controlling these factors would further improve assay performance.

**Figure 8.**
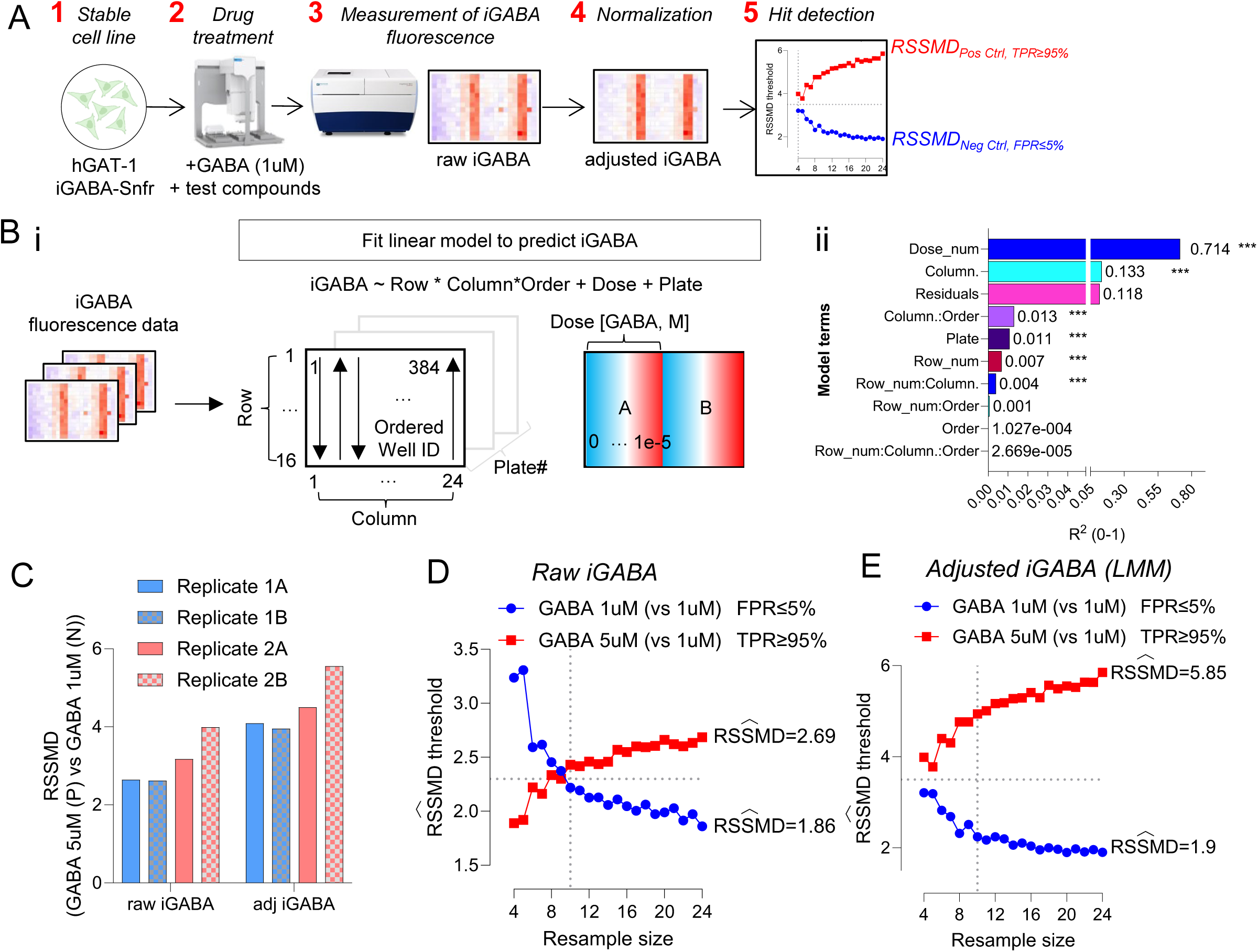
Validation of high-throughput screening strategy to identify GAT-1 agonists using iGABA fluorescence in live cells. (**A**) Proposed work-flow for high-throughput screening includes: 1) stable HeLa cell line expressing iGABA-Snfr and hGAT-1 in 384-well format; 2) application of GABA + test compounds using automated liquid handler (Bravo). The GABA concentration for screening is 1uM, given highly specific GAT-1-dependent uptake at this range. As there are no known GAT-1 agonists, a higher concentration of GABA (5uM) is used as a positive control. Incubation duration = 6-8 hrs; 3) measurement of iGABA fluorescence using automated high-content imaging (IXM); 4) plate normalization and quality control; 5) hit detection and prioritization based on robust strictly standardized mean difference (RSSMD). In a primary screen, compounds whose RSSMD versus on-plate negative controls (GABA 1uM) exceed the RSSMD_FPR ≤ 5%_ threshold are considered hits; compounds whose RSSMD exceed the RSSMD_TPR ≥ 95%_ threshold have effect size similar to positive control. (**B**) A linear model (i) estimates the variance in iGABA explained by covariates (ii). Data are barplots of model R^2^ or residual R^2^ values calculated from ANOVA, and written beside each bar. Statistics are F-tests from ANOVA. ***, *p*<0.001. (**C**) Assay quality based on RSSMD between positive control (GABA 5uM) and negative control (GABA 1uM) for raw iGABA (*left*) and adjusted iGABA (*right*) following plate normalization. Data are RSSMD of iGABA fluorescence (n=28 wells per replicate x condition) from 2 plates, each with 2 replicas (=4 replicates). (**D-E**) Bootstrap simulations from raw iGABA (D) and adjusted iGABA (E) to identify RSSMD screening thesholds for a primary screen without replicates. Thresholds are shown as functions of the number of on-plate replicate controls versus a single test well. The threshold for hit detection (*blue*) maintains FPR≤5%, while the threshold for detecting compounds with effects as large as positive control (GABA 5uM; *red*) maintains TPR≥95%. Vertical dotted lines indicate the RSSMD thresholds @ N=10 replicates. See Method section for further details.

We next compared the performance of adjusting iGABA using the background data extracted from the linear model (LM) or an alternative linear mixed model (LMM) incorporating random effects of well and plate (**Supplementary Fig 4A**). We observed that the LMM adjustment better accounted for well-specific noise (**Supplementary Fig 4B**).

Next we assessed assay quality for the detection of positive modulators of GAT-1. Given the weak effect of 4PBA and no confirmed positive agonist of GAT-1, we chose to model the effects of strong positive modulation by utilizing 5X extracellular GABA (5 µM) as positive control, which should significantly increase iGABA over the test condition (GABA 1uM) based on preliminary data. Initial testing demonstrated that the assay quality index β (derived from robust strictly standardized mean difference (RSSMD) positive vs negative control) was in the *moderate* range (median, 2.91) using raw iGABA and in the *strong* range (median, 4.3) using adjusted iGABA following plate normalization with fixed and random effects from LMM (**Fig 8C**).

To identify the optimal number of control replicates for desired assay performance, we performed bootstrap stimulations for a single concentration primary screen without replicates versus a variable number of on-plate control replicates (4 to 24) for negative (GABA1uM) and positive (GABA 5uM) controls. We observed that a primary screen without replicates could detect significant increases in iGABA above background with a minimum RSSMD threshold = 2.2, assuming 10 negative control replicates. With raw iGABA, the positive control RSSMD threshold was 2.43, assuming 10 positive control replicates, suggesting a narrow window above background to detect compounds whose effect size is less than positive control or to prioritize compounds by their relative effect sizes. However with adjusted iGABA, the positive control RSSMD threshold was 4.94, yielding a larger window for detection and prioritization. This performance window continues to improve with additional replicates using either approach such that with 24 replicates each (positive and negative) the raw iGABA approach has a minimum RSSMD threshold = 1.86 and positive control RSSMD = 2.69, while adjusted iGABA has thresholds 1.90 and 5.85, respectively. These findings suggest the double stable iGABA-Snfr hGAT1 cell line may be suitable for high-throughput compound screening, and that the use of a linear mixed model for plate normalization may improve the prioritization of active compounds.

## 4. Discussion

For patients with epilepsy undergoing genetic testing, identification of a variant in a gene related to epilepsy is generally not sufficient to make a diagnosis, and relies in part on functional assays of gene function. Although mature assays exist for proteins like GAT-1 encoded by *SLC6A1*, the use of radioactivity limits ease of adoption or its application to compound screening. Although recent reports provide extensive functional predictions for over 200 human variants in *SLC6A1*(Silva et al., 2024), there continue to be new human variants needing functional interrogation. Towards those ends, we developed a non-radioactive cell-based assay for functional validation of human GAT-1 coding sequence variants and compound screening using a previously described genetically-encoded GABA fluorescence sensor (iGABA-Snfr) and high-throughput automated microscopy.

### 4.1. Validation of the iGABA assay of GAT-1 function

We established that variant-specific iGABA is highly correlated with GABA uptake estimates from radioactive assays(Silva et al., 2024), and validated the assay’s ability to detect reduced GABA uptake in pathogenic variants and re-classify VUS as pathologic based on reduced iGABA. As predicted, the results of specific GAT-1 variants on the iGABA assay are largely similar to those of [3H]-GABA uptake. Using a conservative pathogenic threshold based on the lower limit (95^th^ CI) of pathogenic variants tested, we demonstrated pathogenic loss-of-function (LoF) in 3 VUS (G111R, S459R, and V511M), corroborating a recent report that showed severe LoF in two (G111R, S495R) and intermediate LoF in another (V511M)(Silva et al., 2024). Meanwhile, 6 additional VUS showed reductions that did not meet our pathogenic threshold (L54F, S574R, F242V, R211C, R419C, and R566H). Based on [3H]-GABA assays, half of these have been previously associated with “typical” GABA uptake (S574R, F242V, R419C), whereas the remaining half have been associated with intermediate LoF (L54F, R211C, and R566H)(Silva et al., 2024). We believe the differences in this case are due to the use of multiple thresholds by other authors, but not due to major discrepancies in the estimated GABA uptake. Together, these observations suggest excellent agreement with prior reports.

We also assessed iGABA as a function of time in a pathogenic variant with residual GABA uptake (V342M) and showed how lower rates of GABA transport are correlated with lower variant-specific steady-state iGABA. Specifically, the change in rate (at least in the case of the V342M variant) can be accounted for mathematically merely by the shift in steady state. This relationship did not have to hold true, as GAT-1 variant-mediated reductions in rate of GABA uptake could in theory reach the same steady state on longer time-scales. However, this does not appear to be the case under these assay conditions (6 hrs), which are likely an overly generous incubation interval compared to the dynamic *in vivo* extracellular environment. Future investigations may explore the extent that this relationship holds true for other patient-derived GAT-1 variants.

### 4.2. Unexpected interactions between WT and mutant GAT-1 variants with residual GABA uptake

We also showed how iGABA responds to the combination of WT with mutant GAT-1, mimicking the heterozygous state that is typical for *SLC6A1*-related disorders. The studies showed the expected additive boost in GAT-1 activity when WT GAT-1 was combined with known pathogenic variants or with VUS with significantly reduced GABA uptake activity (assessed by iGABA), but unexpectedly did not show an additive effect when combined with variants possessing substantial residual GABA uptake activity. It does not appear to be explained by the dynamic range of the assay, given the results from the benign I322M variant. This may suggest a dominant negative effect acting against the compensatory effects of WT GAT-1 supplied *in trans*.

A potential dominant negative (DN) mechanism of SLC6A1-related disease has been raised several times (Carvill et al., 2015; Johannesen et al., 2018; Silva et al., 2024), but to our knowledge most reports to date have been largely consistent with haploinsufficiency. Few reports have convincingly demonstrated a negative interaction between hypomorphic GAT-1 variants and WT GAT-1 function, though relevant data has been reported for only a limited number of variants. Most similar to our own assay, Silva et al. examined combined WT GAT-1 with equimolar variant DNA, including two variants with typical GABA uptake (Asp43Glu, Ile434Met) and five severe LoF variants (Gly63Ser, Tyr140Cys, Ser295L, Leu547Arg, Gly550R), and reported no evidence of DN(Silva et al., 2024). Similar to our data, the authors showed additive increases for variants with severe LoF. However, the authors’ control data suggests that their assay read-out was not in a linear range between 1XWT to 2XWT, making precise interpretation of the results of the variants with typical GABA uptake less clear. Nwosu et al examined a series of GAT-1 variants by [3H]-GABA uptake assay in HEK293T cells with or without 1:1 combinations with WT GAT-1 in the setting of 4PBA treatment(Nwosu et al., 2022), but their method of normalization prevents easy comparison with the results of our study.

Based on *in vivo* studies, the Kang group has stated that the epileptic phenotype of S295L and A288V may be more severe than *SLC6A1* knock-out heterozygotes (Shen et al., 2024), suggesting dominant negativity, but the basis for this interpretation is not clear. By contrast, Lindquist et al(Lindquist et al., 2023) showed similar rates of spike-wave discharges (SWDs) and behavioral arrest in *Slc6a1*-S295L/+ mice compared to a *Slc6a1 +/-* knock-out, though the mouse strains were different (C57BL/6 vs hybrid C57BL/6.129S1, respectively) which may confound a quantitative comparison.

Separately, Shen et al(Shen et al., 2024) reported differences in residual GABA uptake between heterozygous and homozygous mice of the S295L and A288V lines, with similar reductions noted between hets (∼42-51% vs WT), but with greater residual activity (∼30% WT) in A288V/A288V vs S295L/S295L (∼0% WT). This effect is similar to the non-additive effect we observe *in vitro* for other variants, though notably not for the A288V which had virtually no GABA uptake activity in our assay.

Ultimately, we anticipate that it will be important to investigate the potential dominant negative functions of mutant GAT-1 further, as it may have a significant impact on methods of variant interpretation. Although the clinical evidence available suggests a strong correlation between variants with severe LoF in GABA uptake and severe epilepsy phenotypes (e.g. EMAS, EOAE, DEE) (Silva et al., 2024), it has not yet been rigorously established how to interpret variants whose GABA uptake is assessed as falling within a conservative “typical” range (e.g. –49.2% to +47.5% in Silva et al.), which appears to comprise ∼40% of all reported variants. Indeed, although none of the variants in this experiment exceeded our pathogenic threshold (–71.5% reduction vs WT) under “homozygous” conditions, after being combined with WT GAT-1, all of these variants demonstrate GABA uptake similar to that of pathogenic variants combined with WT. This observation raises questions regarding the most appropriate conditions for defining the pathogenicity of mild GAT-1 variants.

Future work should examine the basis for this apparent negative interaction with WT GAT-1 in the “heterozygous” setting, and whether it better correlates with *in vivo* assays of GABA uptake and clinical severity especially for variants with mild loss-of-function.

### 4.3. Case report highlights limitations of GABA uptake assays

We also report the case summary of a boy with NAFE whose M55T variant in *SLC6A1* is associated with normal GABA uptake by the iGABA assay. This variant has been reported once (ClinVar 1307517) but with limited information regarding associated clinical findings or inheritance, and its effects on GABA uptake have not been previously reported. We are confident in our assessment of GABA uptake, given the high correlation between prior radioactive studies and the iGABA assay, as well as the fact that the level of GABA transport activity associated with M55T is indistinguishable from WT, while being higher than any of the pathogenic or VUS variants we studied (15 total, including M55T). This suggests that the M55T variant does not alter GABA transport under these conditions, but we cannot exclude the possibility for different effects under different assay conditions (as discussed in **Section 4.2**), or *in vivo*, nor can we exclude its involvement in other less well-characterized mechanisms that may contribute to *SLC6A1* haploinsufficiency. At the same time, we do not believe that the co-occurring VUS in *GRIK2* is a better explanation, as the patient’s case is not clinically consistent with what is currently known regarding *GRIK2*-associated NDD or epilepsy (see **Section 3.5**). Together, his case highlights the challenge of establishing genetic diagnoses when limited functional evidence is available.

For additional context, we wondered how often clinically ascertained missense variants in *SLC6A1* are not associated with changes in GABA uptake. Based on the most comprehensive analysis of clinically ascertained GAT-1 variants published to date(Silva et al., 2024)(which did not evaluate M55T), there are 28 variants (out of 192 analyzed, ∼14.6%) whose [3H]-GABA transporter activity was unchanged or hyperfunctioning, suggesting it is not a rare phenomenon. Half (14 out of 28) were reported in association with epilepsy. Many of these variants are observed in the general population (gnomAD) and their inheritance patterns not reported, reducing confidence in a causal association. However, at least one nearby variant also located in TM1a, p.F51Y, is not present in population databases, has conflicting interpretations in ClinVar (1 benign, and 1 uncertain), and has been reported to have normal GABA uptake and surface expression (Silva et al., 2024).

Similarly, Silva et al. reported that across epilepsy types, variants reported in association with NAFE (as in our patient) showed GABA uptake that was exclusively in the “typical” or unaffected range. In contrast, variants associated with EMAS were predominantly severe LoF. Although Silva et al. concluded that *SLC6A1* may therefore not be associated with NAFE, this phenomenon could also suggest limitations of heterologous cell lines or assay conditions to recapitulate disease-relevant functions of GAT-1, particularly for variants with mild or no effects on GABA uptake. Ultimately, the pathogenicity of this class of variants warrants further investigation, ideally in animal models where comprehensive evaluations in an intact biological system are possible.

### 4.4 Drug screening

Last we demonstrated the suitability of iGABA for compound screening to identify novel positive modulators of GAT-1. We showed how 4PBA –-a molecular chaperone which appears to increase GAT-1-mediated GABA transport *in vitro* and in preclinical mouse models(Nwosu et al., 2022) – has a mild effect in our assay at specific concentrations of extracellular GABA (1uM). We believe this is largely consistent with the magnitude of effect reported in the literature (∼1.2-1.4x increase), accounting for differences in reported assay conditions(Nwosu et al., 2022). Although 4PBA may show clinical promise, the availability of a high-throughput image-based assay of GAT-1 function may facilitate the identification of more specific positive modulators of GAT-1.

### 4.5 Advantages and limitations of iGABA

Overall, the iGABA assay has the advantages of lower cost and regulatory restrictions compared to radioactive assays, it is portable to any lab with fluorescence microscopy and can be adapted to any high-throughput workflow using automated imaging. We anticipate iGABA will be sensitive to a broad domain of therapeutic agents, including agents that affect uptake rate, surface expression, and intracellular GABA retention, all of which may be biologically relevant for GABA regulation.

We also acknowledge limitations in the iGABA assay. First, the assay measures steady-state intracellular GABA at a single timepoint, not the rate of GABA uptake, making it less sensitive at short incubation durations compared to radioactive uptake, and possibly confounded by other processes affecting the intracellular accumulation of GABA (such as GABA metabolism, sequestration or release). Nevertheless, PAMs enhancing steady-state levels would likely translate to increased GABA clearance *in vivo*, a therapeutic goal for SLC6A1-related disorders like epilepsy. Second, as previously reported(Marvin et al., 2019), compounds with structural similarity to GABA may be a source of false positives in the context of drug screening with iGABA-Snfr, necessitating confirmation of active compounds by orthogonal assays. Third, as discussed, heterologous cell lines may not recapitulate neural-specific modes of GAT-1 regulation and function. Fourth, the described method is limited to coding variants but could be easily extended to assess effects of non-coding or splicing related variants in patient-derived cells, as reported for [3H]-GABA assays(Nwosu et al., 2022).

## 5. Conclusion

In summary, the non-radioactive iGABA assay appears suitable for functional validation of GAT-1 variants as well as high-throughput compound screening to identify positive modulators of GAT-1.

## 6. Data and reagent availability statement

All of the data generated in the present study are available upon request. Plasmids from this study (V006-2, V007-2.2) will be deposited in AddGene. The hGAT1-iGABA cell line is available upon request.

## 7. Acknowledgements

We thank Lee Barrett for technical assistance. This work made use of resources through the F.M. Kirby Assay Development and Screening Facility (ADSF) at Boston Children’s Hospital. We thank Amber Freed (SLC6A1 Connect), Michael Kavanaugh (University of Montana) and Henry Lee (BCH) for helpful discussions, and Jonathan Marvin (Janelia) for providing the data regarding the direct effects of 4PBA on iGABA-Snfr fluorescence.

## 8. Author contributions using the CRediT taxonomy

CM: Conceptualization, Data curation, Formal analysis, Funding acquisition, Investigation, Methodology, Software, Visualization, Supervision, Writing – original draft, Writing – review and editing. GZ: Investigation. GP: Investigation. KW: Writing – review and editing. AP: Funding acquisition, Supervision, Writing – review and editing.

## 13. Supplementary material

Supplementary Figure 1-4 and captions. Supplementary Table 1.

---

**Supplementary Figure 1.**
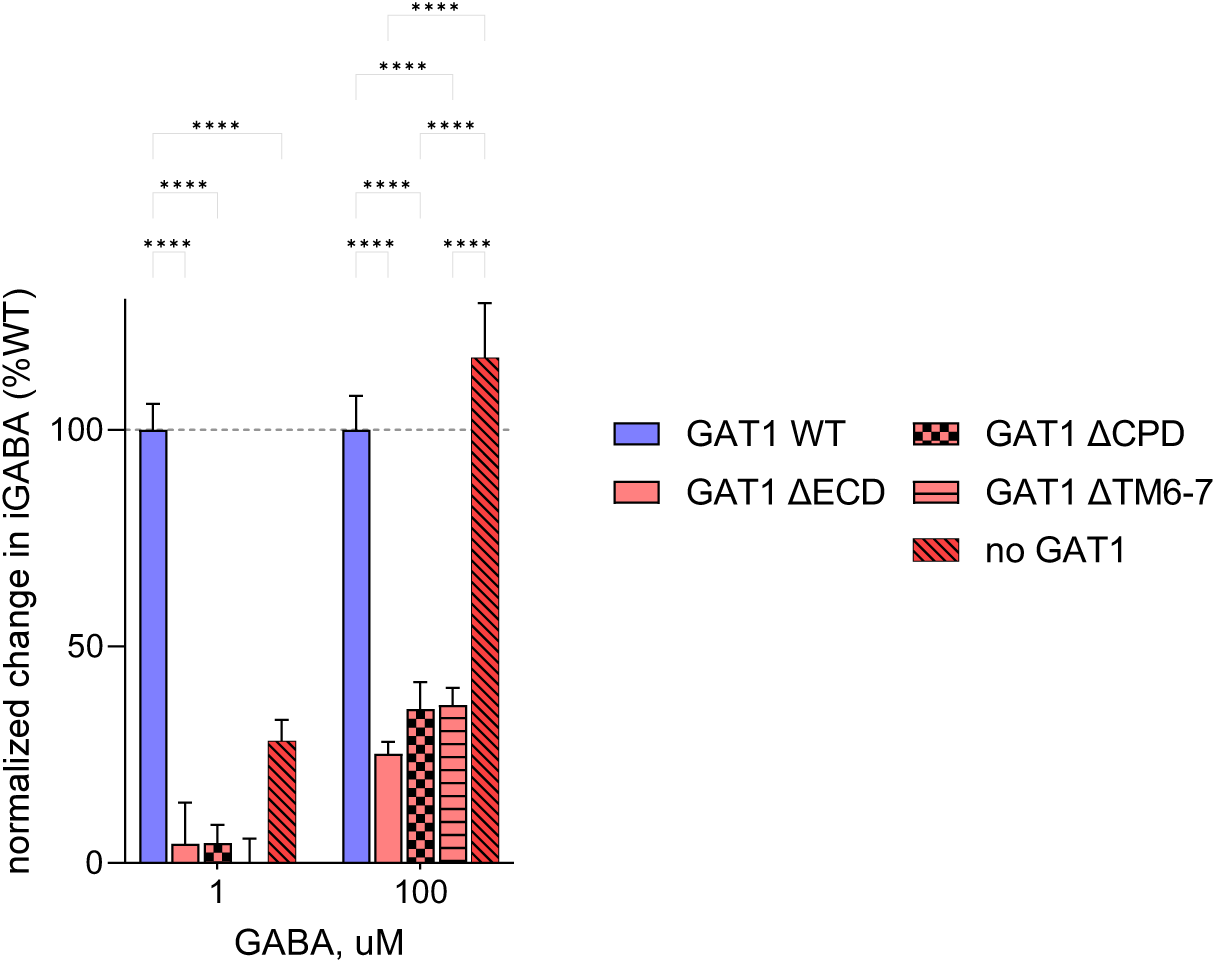
Deletion constructs interfere with non-specific GABA uptake. Transfection of GAT-1 deletion mutants reduces GABA uptake relative to untransfected (“no GAT-1”) control, particularly at higher extracellular GABA concentrations (100uM). Assay format is 96-well plate, transient transfection, incubation time 6-7 hrs. Data are the mean +/-SEM of independent per-well averages of iGABA (4 wells per condition (construct x GABA concentration), 4 sites per well). (Two-way ANOVA, with Tukey’s comparison test. ****, *p*<0.0001)

**Supplementary Figure 2.**
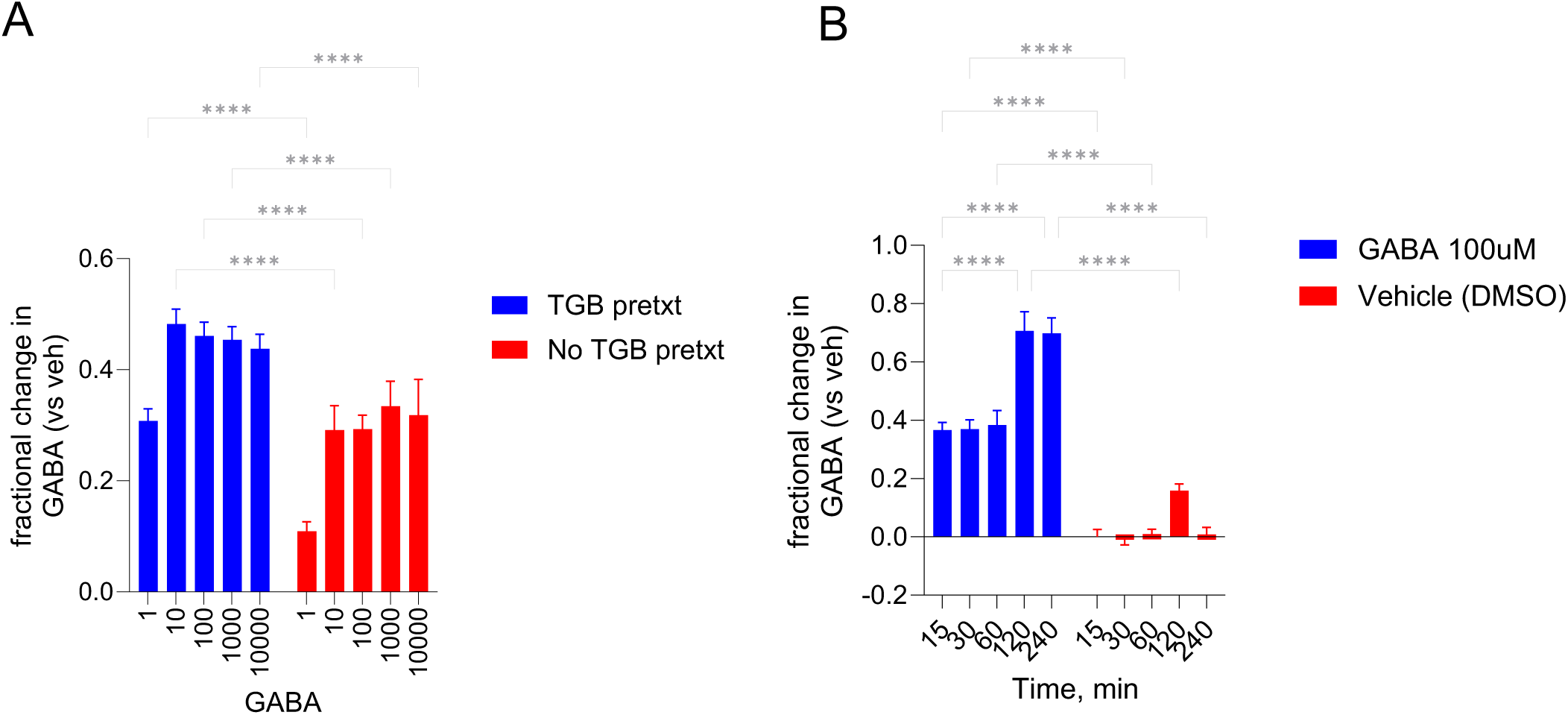
Assay optimization. **A**) Overnight pretreatment with GAT-1 inhibitor (tiagabine) followed by wash-out increases fractional change in iGABA across all GABA concentrations tested. Statistics are ANOVA followed by Sidak’s test (1 family, 5 comparisons, 1 per row (GABA); ****, *p*<0.0001. **B**) GABA fluorescence increases rapidly following incubation with GABA 100uM, but continues to increase at later timepoints (4 sites per well, 45 wells per condition). Statistics are ANOVA followed by Sidak’s test (7 families, 1 comparison per row (GABA), 10 comparisons per column (Time); ****, *p*<0.0001.

**Supplementary Figure 3.**
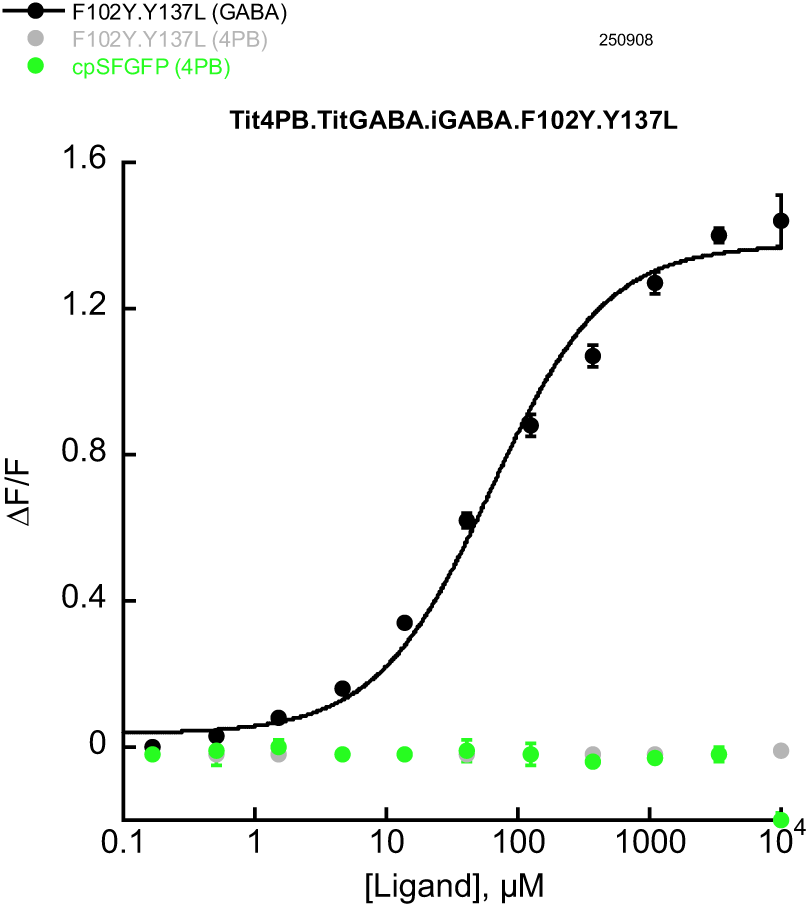
No effect of 4-phenylbutyrate on iGABA-Snfr fluorescence. Data are average normalized fluorescence +/-S.D. (n= 3 replicates per condition) from iGABA-Snfr.F102Y.Y137L incubated with GABA (*black*) or 4-phenylbutyrate (4-PB; *grey*) across a 6-log range of ligand concentrations. Circularly permuted super-folder GFP with 4-PB (*green*) is shown as a control.

**Supplementary Figure 4.**
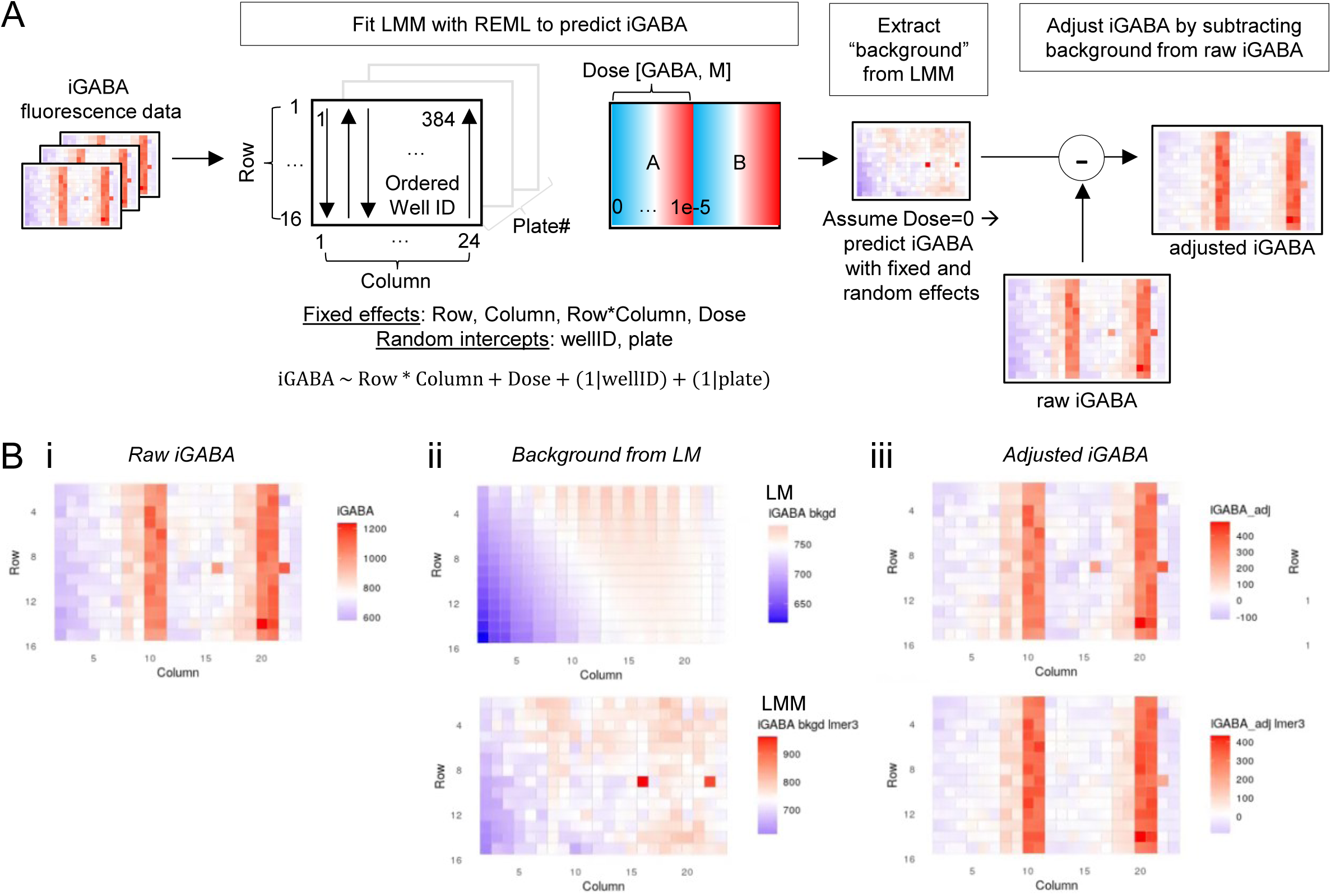
Methods of plate normalization. (**A**) Approach to plate normalization based on linear mixed model (LMM). (**B**) Graphical representation of methods based on linear model vs LMM to extract background (ii) and adjust iGABA (iii).

**Supplementary Table 1.**
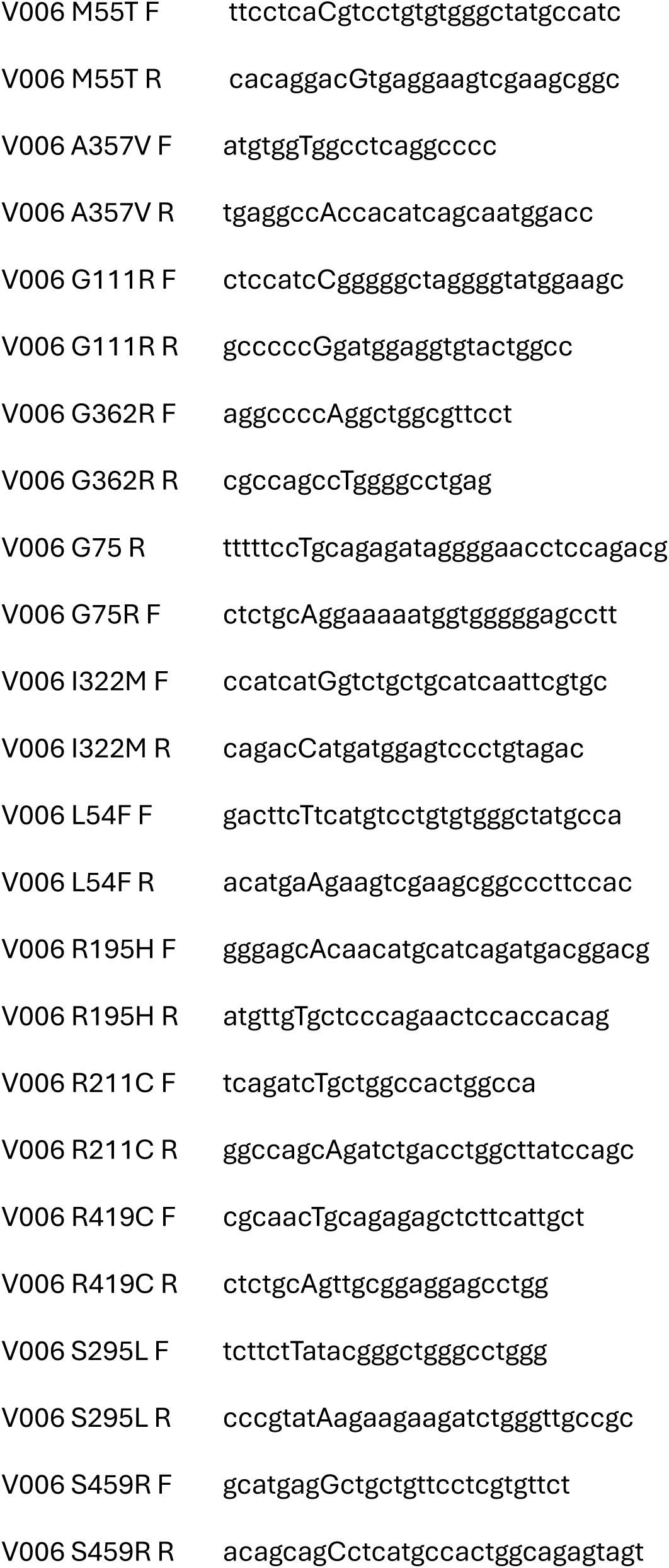
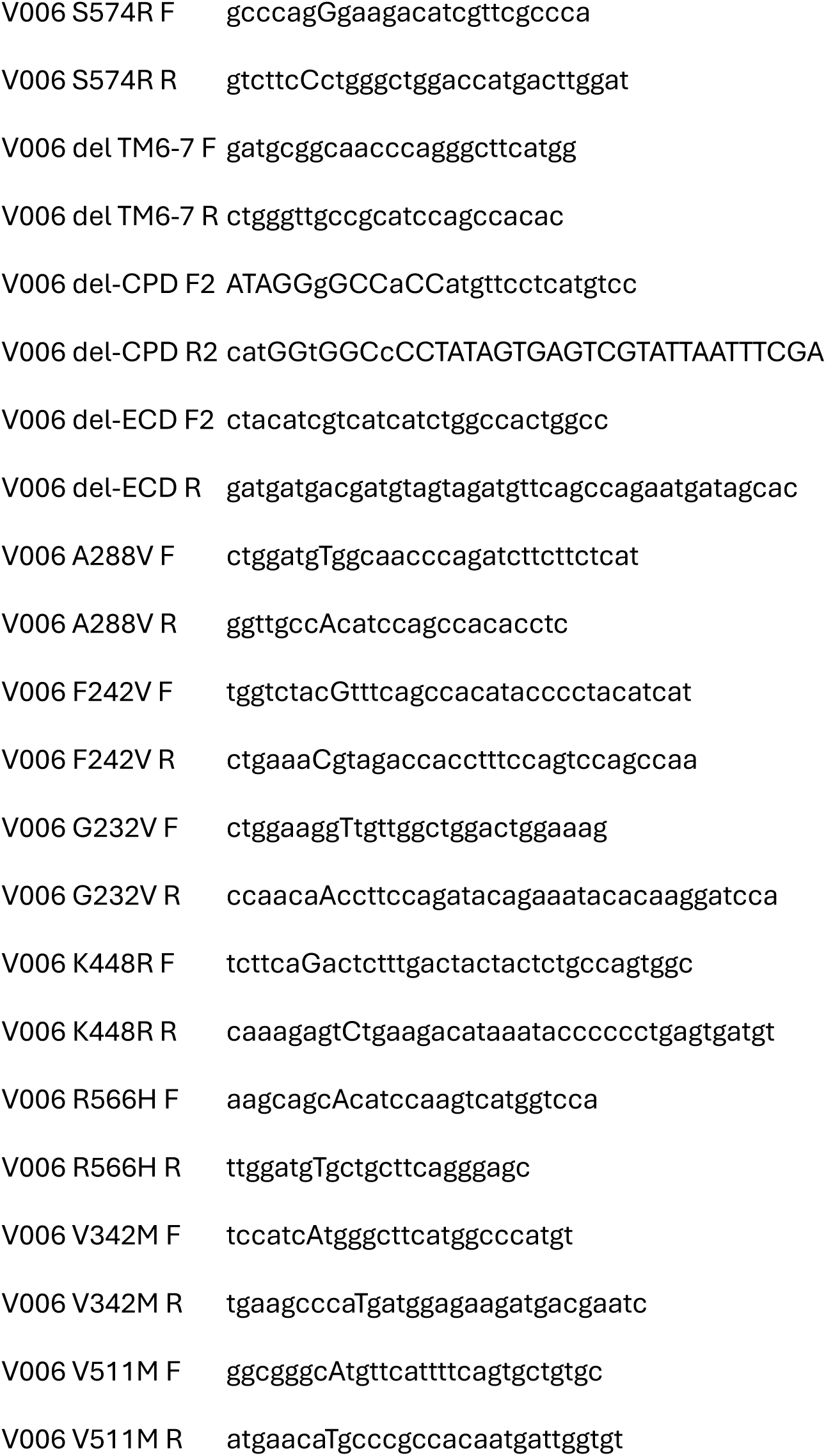
Primers for mutagenesis.

## References

1. Cai, K., Wang, J., Eissman, J., Wang, J., Nwosu, G., Shen, W., Liang, H.-C., Li, X.-J., Zhu, H.-X., Yi, Y.-H., Song, J., Xu, D., Delpire, E., Liao, W.-P., Shi, Y.-W., & Kang, J.-Q. (2019). A missense mutation in SLC6A1 associated with Lennox-Gastaut syndrome impairs GABA transporter 1 protein trafficking and function. Experimental Neurology, 320, 112973. 10.1016/j.expneurol.2019.112973

2. Carvill, G. L., McMahon, J. M., Schneider, A., Zemel, M., Myers, C. T., Saykally, J., Nguyen, J., Robbiano, A., Zara, F., Specchio, N., Mecarelli, O., Smith, R. L., Leventer, R. J., Møller, R. S., Nikanorova, M., Dimova, P., Jordanova, A., Petrou, S., Helbig, I., … Mefford, H. C. (2015). Mutations in the GABA Transporter SLC6A1 Cause Epilepsy with Myoclonic-Atonic Seizures. The American Journal of Human Genetics, 96(5), 808–815. 10.1016/j.ajhg.2015.02.016

3. Eglen, R. M., & Reisine, T. (2009). New Insights into GPCR Function: Implications for HTS. In W. R. Leifert (Ed.), G Protein-Coupled Receptors in Drug Discovery (pp. 1–13). Humana Press. 10.1007/978-1-60327-317-6_1

4. Johannesen, K. M., Gardella, E., Linnankivi, T., Courage, C., Saint Martin, A., Lehesjoki, A.-E., Mignot, C., Afenjar, A., Lesca, G., Abi-Warde, M.-T., Chelly, J., Piton, A., Merritt, J. L., Rodan, L. H., Tan, W.-H., Bird, L. M., Nespeca, M., Gleeson, J. G., Yoo, Y., … Møller, R. S. (2018). Defining the phenotypic spectrum of SLC6A1 mutations. Epilepsia, 59(2), 389–402. 10.1111/epi.13986

5. Lindquist, B. E., Voskobiynyk, Y., Goodspeed, K., & Paz, J. T. (2023). Patient-derived SLC6A1 variant S295L results in an epileptic phenotype similar to haploinsufficient mice. Epilepsia, 64(10), e214–e221. 10.1111/epi.17731

6. Marvin, J. S., Shimoda, Y., Magloire, V., Leite, M., Kawashima, T., Jensen, T. P., Kolb, I., Knott, E. L., Novak, O., Podgorski, K., Leidenheimer, N. J., Rusakov, D. A., Ahrens, M. B., Kullmann, D. M., & Looger, L. L. (2019). A genetically encoded fluorescent sensor for in vivo imaging of GABA. Nature Methods, 16(8), 763–770. 10.1038/s41592-019-0471-2

7. Mattison, K. A., Butler, K. M., Inglis, G. A. S., Dayan, O., Boussidan, H., Bhambhani, V., Philbrook, B., Silva, C., Alexander, J. J., Kanner, B. I., & Escayg, A. (2018). SLC6A1 variants identified in epilepsy patients reduce γ-aminobutyric acid transport. Epilepsia, 59(9), e135–e141. 10.1111/epi.14531

8. Nwosu, G., Mermer, F., Flamm, C., Poliquin, S., Shen, W., Rigsby, K., & Kang, J. Q. (2022). 4-Phenylbutyrate restored γ-aminobutyric acid uptake and reduced seizures in SLC6A1 patient variant-bearing cell and mouse models. Brain Communications, 4(3), fcac144. 10.1093/braincomms/fcac144

9. Rees, E., Han, J., Morgan, J., Carrera, N., Escott-Price, V., Pocklington, A. J., Duffield, M., Hall, L. S., Legge, S. E., Pardiñas, A. F., Richards, A. L., Roth, J., Lezheiko, T., Kondratyev, N., Kaleda, V., Golimbet, V., Parellada, M., González-Peñas, J., … Owen, M. J. (2020). De novo mutations identified by exome sequencing implicate rare missense variants in SLC6A1 in schizophrenia. Nature Neuroscience, 23(2), 179–184. 10.1038/s41593-019-0565-2

10. Richerson, G. B., & Wu, Y. (2004). Role of the GABA Transporter in Epilepsy. In D. K. Binder & H. E. Scharfman (Eds.), Recent Advances in Epilepsy Research (Vol. 548, pp. 76–91). Springer US. 10.1007/978-1-4757-6376-8_6

11. Shen, W., Nwosu, G., Honer, M., Clasadonte, J., Schmalzbauer, S., Biven, M., Langer, K., Flamm, C., Poliquin, S., Mermer, F., Dedeurwaerdere, S., Hernandez, M.-C., & Kang, J.-Q. (2024). γ-Aminobutyric acid transporter and GABAA receptor mechanisms in Slc6a1+/A288V and Slc6a1+/S295L mice associated with developmental and epileptic encephalopathies. Brain Communications, 6(2), fcae110. 10.1093/braincomms/fcae110

12. Silva, D. B., Trinidad, M., Ljungdahl, A., Revalde, J. L., Berguig, G. Y., Wallace, W., Patrick, C. S., Bomba, L., Arkin, M., Dong, S., Estrada, K., Hutchinson, K., LeBowitz, J. H., Schlessinger, A., Johannesen, K. M., Møller, R. S., Giacomini, K. M., Froelich, S., Sanders, S. J., & Wuster, A. (2024). Haploinsufficiency underlies the neurodevelopmental consequences of SLC6A1 variants. The American Journal of Human Genetics, 111(6), 1222–1238. 10.1016/j.ajhg.2024.04.021

13. Stolz, J. R., Foote, K. M., Veenstra-Knol, H. E., Pfundt, R., ten Broeke, S. W., de Leeuw, N., Roht, L., Pajusalu, S., Part, R., Rebane, I., Õunap, K., Stark, Z., Kirk, E. P., Lawson, J. A., Lunke, S., Christodoulou, J., Louie, R. J., Rogers, R. C., Davis, J. M., … Swanson, G. T. (2021). Clustered mutations in the GRIK2 kainate receptor subunit gene underlie diverse neurodevelopmental disorders. American Journal of Human Genetics, 108(9), 1692–1709. 10.1016/j.ajhg.2021.07.007

14. Wu, Y., Wang, W., Díez-Sampedro, A., & Richerson, G. B. (2007). Nonvesicular inhibitory neurotransmission via reversal of the GABA transporter GAT-1. Neuron, 56(5), 851–865. 10.1016/j.neuron.2007.10.021

15. Zhang, X. D. (2011). Illustration of SSMD, z score, SSMD*, z* score, and t statistic for hit selection in RNAi high-throughput screens. Journal of Biomolecular Screening, 16(7), 775–785. 10.1177/1087057111405851

